# S100A9-Dependent CXCR2^hi^ Neutrophils Mediate Systemic Immune Suppression and Checkpoint Resistance in Metastatic TNBC

**DOI:** 10.64898/2026.06.09.731132

**Authors:** Fulya Koksalar Alkan, Ahmet Burak Caglayan, Hilmi K. Alkan, Eunmi Lee, Raziye Piranlioglu, Cordarryl Jones, Max Alimadadi, Elayne Benson, Alicia Arnold, Adriana Langer Gramer, Thomas Vogl, Greg Dyson, Ahmed Chadli, Mustafa Guzel, Sabine Kasimir-Bauer, Hadeel Assad, Julie Boerner, Morhaf Al-Achkar, Asfar S. Azmi, Nouri Neamati, Gurkan Ozturk, Roni Bollag, Catherine C. Hedrick, Max S. Wicha, Huidong Shi, Hasan Korkaya

## Abstract

Most high-dimensional studies of tumor-immune interactions focus on metastatic models, limiting insight into how immune remodeling in primary tumors shapes metastatic competence. Here, integrating single-cell RNA sequencing, CyTOF, and functional studies across metastatic (4T1) and non-invasive (EMT6) triple-negative breast cancer (TNBC) murine models, we define tumor state-specific immune programs that distinguish metastatic competence. Tumors with metastatic capacity uniquely drive early bone marrow expansion of CXCR2⁺ neutrophils, which infiltrate primary tumors acquiring a CXCL2-producing phenotype that promotes EMT-associated cancer stem cell (CSC) plasticity. This program depends on TGF-β/CEBPD-mediated induction of S100A9. Elevated CXCL2, together with G-CSF, establishes a feed-forward circuit that drives systemic neutrophil mobilization and recruitment to distant organs, where neutrophil-derived S100A8/A9 (calprotectin) promotes MET-driven CSC outgrowth and metastatic colonization. Clinically, gene signatures associated with CXCR2⁺ neutrophils predict poor survival in TNBC patients, whereas monocyte/macrophage (CX3CR1⁺) and T cell activation signatures correlate with improved outcomes. S100A9 ablation disrupts this cascade and enhances immunotherapy responsiveness, defining a TGF-β/S100A9/CXCR2 axis linking immune remodeling, CSC plasticity and metastasis.

**Highlights:**

- Metastatic TNBC engages a TGF-β/C/EBPδ/S100A9 axis that expands CXCR2⁺ neutrophils
- Non-invasive EMT6 tumors retain a CX3CR1⁺ monocyte/macrophage and T-cell landscape
- CXCR2^+^ neutrophils in pre-metastatic niches suppress T cell response while promoting tumor cell proliferation
- S100A9 loss redirects myelopoiesis and potentiates anti-PD-L1 in TNBC models

**In Brief:** Alkan et al. dissect how tumor state programs the myeloid compartment in TNBC. Metastatic 4T1 tumors uniquely engage a TGF-β/C/EBPδ/S100A9 axis driving CXCR2⁺ neutrophil expansion and CXCL2/G-CSF-dependent systemic mobilization, coupling immune remodeling to EMT/MET cancer-stem-cell plasticity, while S100A9 loss restores CX3CR1⁺ myeloid identity and unlocks checkpoint-inhibitor responsiveness.

## Introduction

Triple-negative breast cancer (TNBC) is an aggressive disease characterized by early metastatic dissemination, poor clinical outcomes, and limited response to immunotherapy in advanced disease^1,2^. A defining feature of TNBC progression is the establishment of an immunosuppressive tumor microenvironment (TME) enriched with heterogeneous myeloid populations, including tumor-associated neutrophils (TANs) and myeloid-derived suppressor cells (MDSCs), which restrain anti-tumor immunity and promote therapeutic resistance^3–7^. Although recent advances using single-cell RNA sequencing (scRNA-seq) and mass cytometry (CyTOF) have helped define the cellular architecture of the TME, these studies have largely focused on metastatic disease, limiting insight into how tumor-intrinsic states shape local and systemic immune remodeling. Emerging evidence indicates that tumors can systemically reprogram hematopoiesis, particularly through expansion of granulocytic lineages resulting in elevated neutrophil-to-lymphocyte ratios, which are associated with poor prognosis^8–10^. However, the mechanisms linking these systemic alterations to metastatic competence remain incompletely defined.

Comparative tumor models provide a unique framework to address this gap. The syngeneic 4T1 model recapitulates key features of human metastatic TNBC, including robust dissemination and establishment of systemic immune suppression, whereas the EMT6 model represents a non-invasive state capable of eliciting protective anti-tumor immunity following tumor resection^11–13^. Utilizing these TNBC models, we interrogated how tumor state governs immune reprogramming across bone marrow, primary tumors, and distant organs. We find that metastatic tumors uniquely drive early expansion of CXCR2⁺ neutrophil progenitors in the bone marrow promoting their mobilization via G-CSF-dependent signaling^14^, consistent with prior reports linking tumor-derived G-CSF to granulopoiesis and pre-metastatic niche formation^9^. Upon tumor infiltration, these neutrophils undergo transcriptional reprogramming characterized by TGF-β/CEBPD-mediated induction of S100A9, a key regulator of inflammatory myeloid function and MDSC activity^7,15^. This reprogramming licenses a CXCL2-producing phenotype, establishing a feed-forward chemokine circuit that amplifies recruitment and retention of CXCR2⁺ neutrophils while promoting epithelial-mesenchymal transition (EMT)-associated cancer stem cell (CSC) plasticity, processes previously linked to inflammatory cytokine signaling including IL-1β and CXCR2 ligands^6,8^.

Systemically, this G-CSF/CXCL2/S100A9 axis drives BM expansion/mobilization and trafficking of immature Ly6G⁺Ly6C⁻CXCR2⁺ neutrophils to pre-metastatic niches, where neutrophil-derived S100A8/A9 (calprotectin) conditions an immunosuppressive microenvironment characterized by myeloid skewing, T cell dysfunction, and impaired antigen presentation^5,7^. In this context, calprotectin has been implicated in promoting tumor progression and metastatic niche formation through modulation of inflammatory signaling and recruitment of suppressive myeloid populations^6,15^. Consistent with these mechanisms, we have previously demonstrated that systemic expansion of granulocytic MDSCs is required for metastatic outgrowth in TNBC models^11,12^. In contrast, non-invasive tumors fail to engage this myeloid reprogramming cascade and instead support lymphocyte expansion and concomitant anti-tumor immunity^11^.

Here, we demonstrate that genetic ablation of S100A9 disrupts this feed-forward circuit at multiple levels, impairing bone marrow expansion and CXCR2-dependent mobilization of neutrophils, limiting their tumor infiltration and CXCL2 production, and preventing the establishment of an immunosuppressive pre-metastatic niche. Loss of S100A9 further attenuates neutrophil functional polarization, resulting in reduced calprotectin-mediated signaling, restoration of T cell activity, and enhanced antigen presentation in distant tissues. Functionally, these changes translate into diminished metastatic colonization and a marked increase in sensitivity to immune checkpoint blockade. Notably, these mechanistic insights are supported by clinical analyses demonstrating that gene signatures associated with CXCR2⁺ neutrophils (including S100A8/A9, CXCL2, and CSF3R) correlate with poor survival in TNBC patients, whereas signatures linked to CX3CR1⁺ monocyte/macrophage populations and T cell activation associate with improved outcomes. Collectively, these findings define a metastasis-specific TGF-β/S100A9/CXCR2 axis that integrates tumor-driven myelopoiesis, neutrophil reprogramming, and cancer stem cell plasticity, and identify calprotectin-dependent neutrophil signaling as a tractable therapeutic vulnerability to disrupt immune-mediated metastatic progression in TNBC.

## RESULTS

### Metastatic murine TNBC model identifies neutrophil-dominated immune remodeling which is associated with poor clinical outcome

To define early immune programs associated with metastatic competence, we performed integrated single-cell RNA sequencing and translational analyses comparing metastatic 4T1 and non-invasive EMT6 TNBC models. To minimize inter-tumor variability, five independent tumors per model were pooled prior to analysis (Figure S1A).

Single-cell profiling revealed that 4T1 and EMT6 tumors establish fundamentally distinct immune ecosystems. 4T1 tumors^11,12^ were markedly enriched for myeloid populations, particularly neutrophils and monocytes, whereas EMT6 tumors exhibited increased lymphoid infiltration (Figures 1A-1C). High-resolution subclustering further resolved multiple granulocytic and monocytic populations, CXCR2^hi^ tumor associated neutrophils (TAN), Ccl3hi TANs, and macrophages. Notably, 4T1 tumors displayed a pronounced expansion of distinct neutrophil states, including immature, inflammatory, and a transcriptionally defined CXCR2^hi^ TAN subpopulation, which was largely absent or minimally represented in EMT6 tumors (Figure 1D). Hierarchical clustering confirmed segregation of myeloid and lymphoid transcriptional programs, with neutrophils forming a distinct inflammatory cluster (Figure S2).

**Figure 1.**
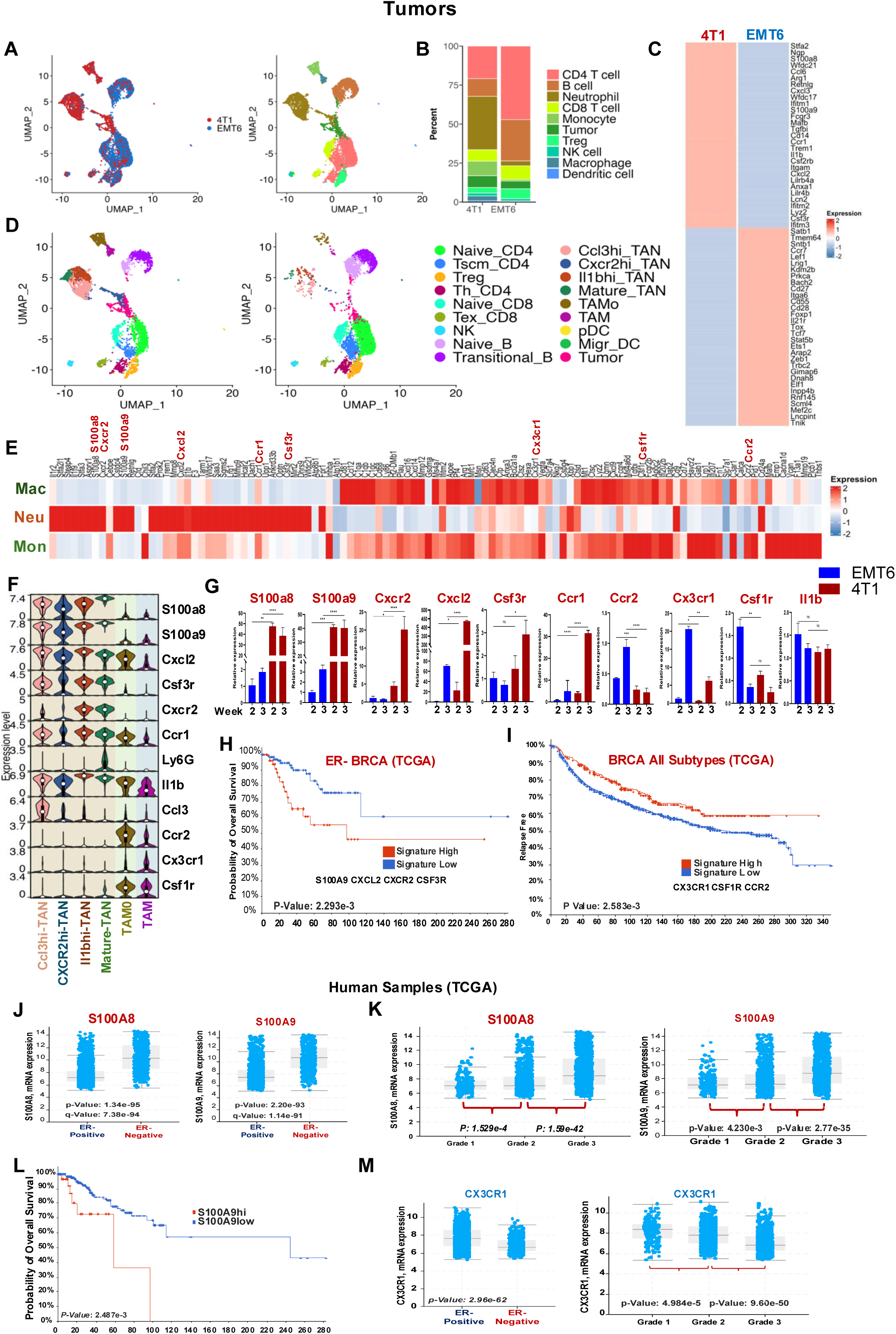
Myeloid-driven immune remodeling distinguishes metastatic and non-invasive TNBC and predicts clinical outcome. **(A)** UMAP visualization of single-cell RNA sequencing data from pooled primary tumors (five independent tumors per model) showing segregation of cells from metastatic 4T1 and non-invasive EMT6 tumors. **(B)** UMAP colored by annotated cell types, identifying major immune and tumor cell populations across conditions. **(C)** Relative proportions of immune cell populations in 4T1 and EMT6 tumors, demonstrating enrichment of myeloid populations, particularly neutrophils and monocytes, in 4T1 tumors and increased lymphoid populations in EMT6 tumors. **(D)** High-resolution subclustering of tumor-infiltrating immune cells, identifying granulocytic and monocytic subsets, including CXCR2hi tumor associated neutrophils (TAN), Ccl3hi TANs, and macrophage populations. **(E)** Heatmap of differentially expressed genes across myeloid populations, highlighting enrichment of neutrophil-associated transcriptional programs in 4T1 tumors compared to EMT6 tumors. **(F)** Violin plots showing expression of key lineage and functional markers (*S100a8, S100a9, Cxcl2, Csf3r, Cxcr2, Ccr1, Ly6g, Il1b, Csf1r, Cx3cr1*) across identified myeloid subsets. **(G)** qRT-PCR based quantification of selected neutrophil– and myeloid-associated gene expression across tumor models over time, demonstrating increased expression of inflammatory and granulocytic markers in 4T1 tumors. **(H, I)** Kaplan-Meier survival analysis of breast cancer patients (TCGA datasets) stratified by neutrophil-associated gene signatures (*S100A8, S100A9, CXCL2, CXCR2, CSF3R*) or monocyte/macrophage-associated signatures (*CX3CR1, CSF1R, CCR2*), showing association of neutrophil signatures with poor survival and monocyte/macrophage signatures with improved survival. **(K, L)** Analysis of human breast cancer datasets (TCGA) showing increased expression of *S100A8* and *S100A9* in aggressive tumor subtypes and higher histological grades. **(M)** Kaplan-Meier survival analysis stratified by *S100A9* expression, demonstrating association of high expression with reduced overall survival. **(N)** Expression of *CX3CR1* across breast cancer subtypes and grades, showing association with less aggressive disease features. Data represent mean ± SEM; statistical significance determined Welch-t-test (*p **<**0.05, **p **<**0.01, *** p **<** 0.005, **** p **<** 0.001)

Differential gene expression (DGE) analysis demonstrated selective upregulation of neutrophil-associated genes, including *S100a8*, *S100a9*, *Cxcr2*, *Cxcl2*, and *Csf3r*, in 4T1 tumors, whereas EMT6 tumors were enriched for antigen presentation and lymphocyte activation programs (Figures 1E-1G and Table S1A). Importantly, the CXCR2^hi^ TAN population exhibited particularly high expression of S100A8/A9 and CXCL2 signaling components (Tabel S1B), suggesting a specialized, pro-inflammatory and tumor-promoting neutrophil state within metastatic tumors.

These tumor-specific immune programs were clinically relevant. CXCR2⁺ neutrophil-associated gene signatures predicted significantly worse survival in TNBC patients, whereas monocyte/macrophage-associated signatures (e.g., *CX3CR1*) correlated with improved outcomes in breast cancer patients across all subtypes (Figures 1H-1I). Consistent with this, TCGA analyses showed that *S100A8/A9* expression in TNBC correlates with higher tumor grade and poor survival, whereas *CX3CR1* expression is significantly lower in TNBC and reduced with higher grade (Figures 1J-1M).

Together, these findings establish that metastatic TNBC is characterized by a neutrophil-dominated immune landscape driven in part by expansion of a CXCR2^hi^ TAN subpopulation, which is associated with inflammatory signaling, tumor progression, and adverse clinical outcomes. In contrast, non-invasive tumors maintain a more immune-activated microenvironment enriched for lymphoid and monocyte/macrophage populations linked to favorable outcome.

### Non-invasive EMT6 tumor model sustains cytotoxic T cell immunity which is associated with improved patient survival

Given the pronounced myeloid skewing observed in metastasis-competent tumors, we next examined lymphoid compartments across tumor and systemic sites. Single-cell transcriptional profiling revealed that EMT6 tumors are enriched for CD4⁺ and CD8⁺ T cell populations, whereas 4T1 tumors exhibit reduced lymphocyte representation and altered T cell states (Figures 2A-2B).

**Figure 2.**
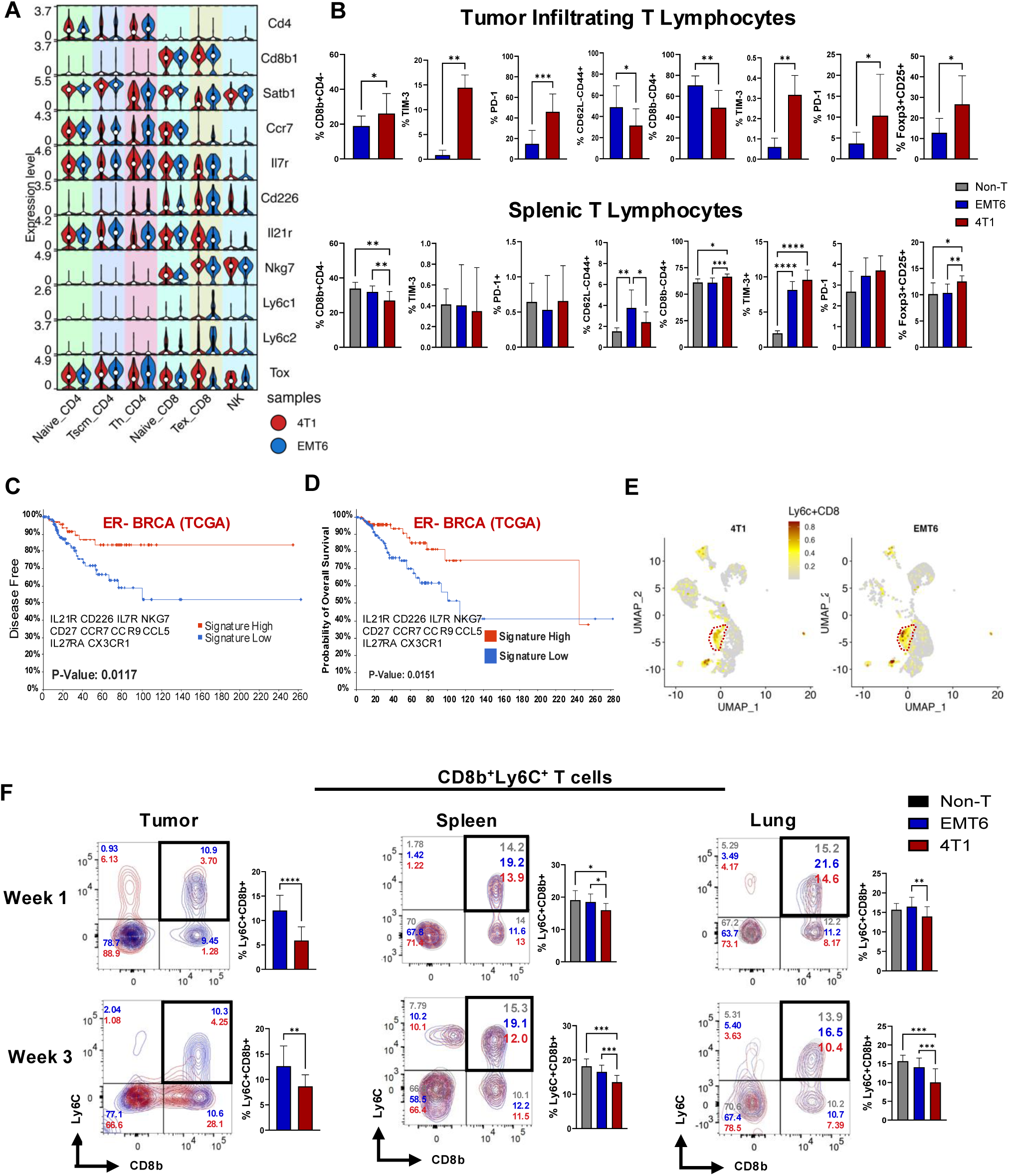
Non-invasive EMT6 tumors sustain cytotoxic T immunity locally and systemically and are associated with favorable clinical outcomes. **(A)** Violin plots showing expression of T cell-associated genes (*Cd4, Cd8b1, Satb1, Ccr7, Il7r, Cd226, Il21r, Nkg7, Ly6c1, Ly6c2, Tox*) across CD4⁺ T cells, CD8⁺ T cells, and NK cells in 4T1 and EMT6 tumors, highlighting enhanced activation and effector programs in EMT6-derived lymphoid populations. **(B)** Flow cytometry analyses of tumor-infiltrating (top) and splenic (bottom) T cell populations and activation/exhaustion markers, including CD8⁺ T cells, CD4⁺ T cells, PD-1, TIM-3, and FoxP3⁺CD25⁺ regulatory T cells, demonstrating increased cytotoxic T cell abundance and reduced exhaustion in EMT6 tumors and spleens compared to 4T1 tumors and spleens (n=10 each). **(C-D)** Kaplan-Meier analyses of TCGA ER⁻ breast cancer cohorts showing improved disease-free survival (C) and overall survival (D) in patients with high T cell activation gene signatures. **(E)** UMAP projection of tumor-infiltrating immune cells highlighting the spatial distribution and enrichment of CD8⁺Ly6C⁺ T cells in EMT6 compared to 4T1 tumors. **(F)** Flow cytometry analysis of T cell populations across tumor, spleen, and lung compartments at early (week 1) and later (week 3) time points in non-tumor-bearing, EMT6, and 4T1 mice. Quantification demonstrates sustained enrichment of cytotoxic CD8⁺Ly6C⁺ T cells in EMT6 across local and systemic compartments, whereas 4T1 tumors exhibit reduced frequencies consistent with impaired cytotoxic T cell immunity. One-Way ANOVA mean ± SEM test applied to compare tumor-derived values and Welch-t-test applied to compare spleen and lungs-derived values *p **<**0.05, **p **<**0.01, ***p **<**0.005, **** p **<** 0.001

At the transcriptional level, T cells in EMT6 tumors displayed elevated expression of activation-and memory-associated genes, including *Cd8b1, Ccr7, Il7r, Cd226,* and *Nkg7*, consistent with functional anti-tumor immunity. In contrast, 4T1 tumors exhibited features of T cell dysfunction, including increased expression of exhaustion-associated markers and enrichment of regulatory T cell populations (Figures 2A-2B).

Importantly, these differences extended beyond the tumor microenvironment. Flow cytometric analysis demonstrated that EMT6-bearing mice maintain higher frequencies of functional CD8⁺ T cells both within tumors and in peripheral compartments, including spleen and lung, whereas 4T1 tumors are associated with reduced cytotoxic T cell abundance and increased immunoregulatory populations (Figure 2B, 2F). Notably, CD8⁺Ly6C⁺ T cells, previously implicated in potent cytotoxic and effector functions^11^, were significantly enriched in EMT6 tumors as well as in systemic compartments, including spleen and lung, particularly at early time points (week 1), and remained elevated relative to 4T1 at later stages (week 3) (Figures 2E-2F).

Consistent with these observations, gene signatures associated with T cell activation and effector function were strongly associated with improved disease-free and overall survival in ER⁻ breast cancer cohorts (Figures 2C-2D), supporting the clinical relevance of sustained cytotoxic T cell immunity.

Together, these findings demonstrate that non-invasive EMT6 tumors preserve both local and systemic cytotoxic T cell responses, whereas metastatic 4T1 tumors promote T cell dysfunction and immune suppression. This divergence in lymphoid programming complements the neutrophil-dominated myeloid landscape observed in metastatic disease and highlights a coordinated reprogramming of both innate and adaptive immunity during early tumor progression.

### TGF-β/S100A9 signaling drives expansion of CXCR2⁺CXCL2⁺ tumor neutrophils and dictates myeloid fate

We next investigated the mechanisms underlying neutrophil expansion in metastasis competent tumors. UMAP analysis revealed enrichment of CXCR2⁺, CXCL2⁺, and S100A8/A9⁺ myeloid populations in 4T1 tumors, consistent with activation of a granulocytic inflammatory program (Figure 3A). In contrast, EMT6 tumors displayed a relative enrichment of a monocyte/macrophage population characterized by CX3CR1 expression, indicative of a more differentiated, tissue resident and immunologically permissive myeloid compartment. Flow cytometry further confirmed the early and preferential expansion of CD11b⁺CXCR2⁺ neutrophils and CXCR2⁺CXCL2⁺ subsets in 4T1 tumors, whereas EMT6 tumors maintained a lower frequency of these granulocytic populations and instead favored CX3CR1 monocyte/macrophage lineages (Figures 3B-3C).

**Figure 3.**
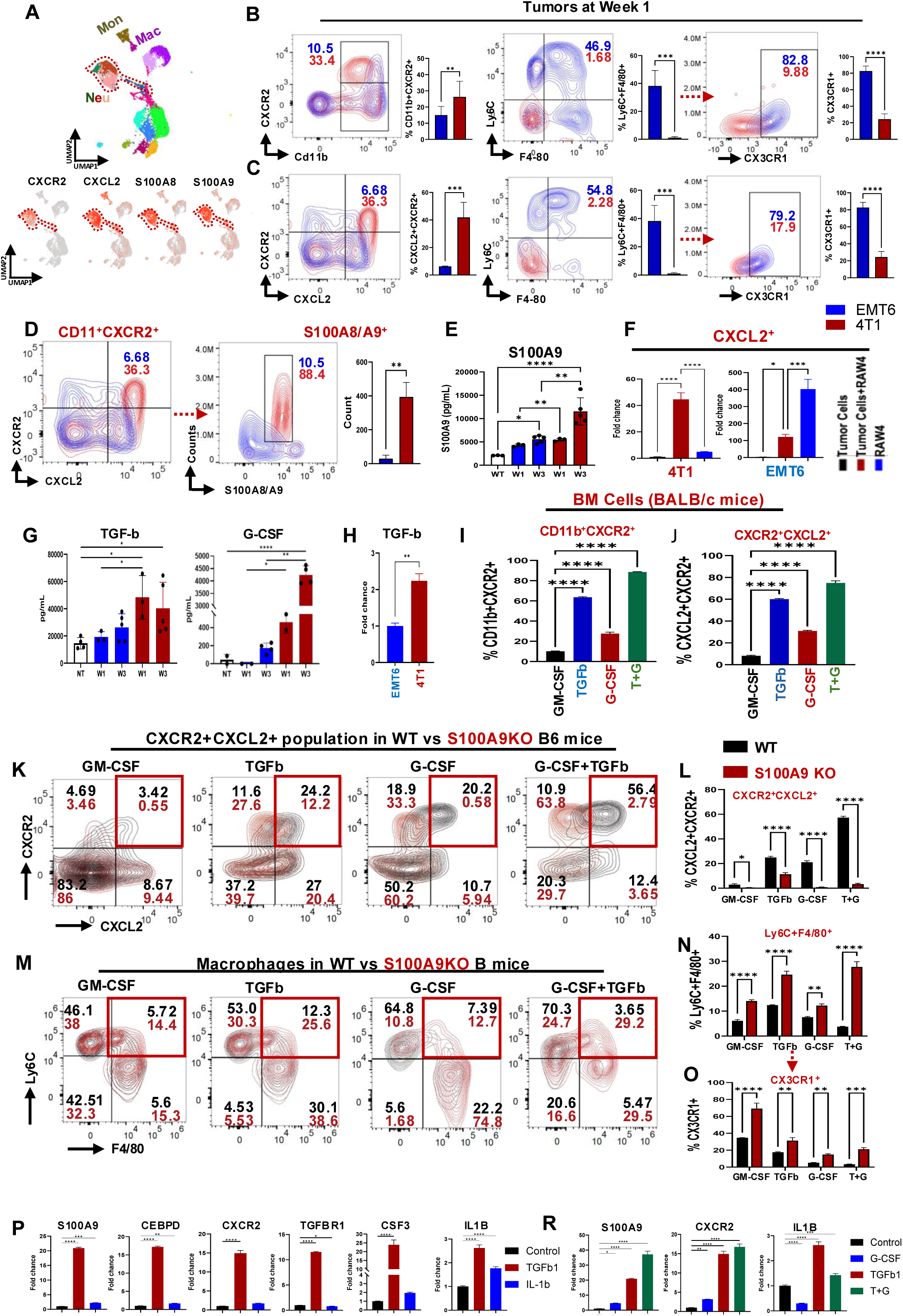
TGF-β-S100A9 signaling drives expansion of CXCR2⁺CXCL2⁺ neutrophils and dictates myeloid lineage fate. (**A**) UMAP visualization of tumor-infiltrating myeloid populations highlighting enrichment of neutrophils (Neu), monocytes (Mon), and macrophages (Mac), with feature plots showing expression of *CXCR2, CXCL2, S100A8,* and *S100A9* in neutrophil-enriched clusters in 4T1 tumor. **(B-C)** Flow cytometric analysis of tumors at week 1 demonstrating early expansion of CD11b⁺CXCR2⁺ and CXCR2⁺CXCL2⁺ neutrophil populations in 4T1 compared to EMT6 tumors (n=10 each). Representative contour plots and quantification are shown. (Welch-t-test applied to compare tumor-derived values **p **<**0.01, ***p **<**0.005, **** p **<** 0.001) **(D)** Quantification of CD11b⁺CXCR2⁺ and S100A8/A9⁺ neutrophil populations, demonstrating significant enrichment of inflammatory neutrophils in 4T1 tumors (n=5 each). (Welch-t-test applied to compare tumor-derived values **p **<**0.01) **(E)** ELISA-based quantification of S100A9 levels in tumor lysates across time points, showing progressive elevation in metastatic 4T1 tumors. **(F)** Quantification of *CXCL2* expression in RAW264.7 myeloid cells that are co-cultured with tumor cells, demonstrating increased induction in response to 4T1 compared to EMT6 tumor cells (One-Way ANOVA mean ± SEM test applied compared to 4T1 or EMT6 values *p **<**0.05, ***p **<**0.005, **** p **<** 0.001). **(G)** Quantification of serum derived circulating TGF-β and G-CSF levels in tumor-bearing mice over time by ELISA One-Way ANOVA mean ± SEM test applied compared to WT/Non-T values **(H)** RT-PCR based relative expression measurement of TGF-â signaling components in tumor models at the early stage (n=5 each) (Welch-t-test applied to compare tumors **p **<**0.01). *p **<**0.05, **** p **<** 0.001, Welch-t-test applied compared between weeks **p **<**0.01). **(I, J)** Flow cytometric analysis of Balb/c bone marrow cells showing induction of CD11b⁺CXCR2⁺ and CXCR2⁺CXCL2⁺ populations in response to GM-CSF, TGF-β, and G-CSF stimulation (n=2, One-Way ANOVA mean ± SEM test applied compared to GM-CSF *p **<**0.05, ***p **<**0.005, **** p **<** 0.001).). **(K)** Representative Quantification of CXCR2⁺CXCL2⁺ neutrophil populations under indicated cytokine conditions in C57BL6 WT and S100A9 KO mice. **(L)** Analysis of CXCR2⁺CXCL2⁺ populations in WT versus S100A9-deficient bone marrow cells, demonstrating impaired expansion in S100A9-deficient conditions (n=3 biological replicates of each) (Welch-t-test applied for the comparison within conditions *p **<**0.05, ***p **<**0.005, **** p **<** 0.001). **(M, N)** Flow cytometric analysis of macrophage differentiation (Ly6C⁻F4/80⁺) in wild-type and S100A9-deficient bone marrow cultures under cytokine stimulation (n=3 biological replicates of each) (Welch-t-test applied for the comparison within conditions *p **<**0.05, ***p **<**0.005, **** p **<** 0.001). **(O)** Quantification of CX3CR1⁺ monocyte/macrophage populations under indicated conditions n=3 biological replicates of each) (Welch-t-test applied for the comparison within conditions *p **<**0.05, ***p **<**0.005, **** p **<** 0.001). **(P, R)** Gene expression analysis of wild-type and S100A9-deficient bone marrow cultures by RT-PCR showing induction of S100A9, CEBPD, CXCR2, TGFBR1, CSF3, and IL1B in response to TGF-β and G-CSF signaling (n=2, biological replicates) (One-Way ANOVA mean ± SEM test applied compared to GM-CSF *p **<**0.05, ***p **<**0.005, **** p **<** 0.001).

These cells exhibited elevated *S100A8/A9* expression (Figure 3D), which was also increased systemically (Figure 3E), indicating tumor-driven inflammatory reprogramming. Co-culture experiments demonstrated that 4T1 tumor cells directly induce CXCL2 production in myeloid cells (Figure 3F), suggesting active tumor-immune instruction.

Upstream, 4T1 tumors exhibited elevated TGF-β and G-CSF levels early after implantation (Figures 3G-3H). These cytokines synergistically promoted generation of CXCR2⁺CXCL2⁺ neutrophils from bone marrow precursors (Figures 3I-3J), establishing a feed-forward granulopoietic axis.

Importantly, this process was critically dependent on S100A9. Stimulation of bone marrow cells with TGF-β or G-CSF alone induced modest expansion of CD11b⁺CXCR2⁺ and CXCR2⁺CXCL2⁺ populations; however, combined TGF-β and G-CSF treatment resulted in a markedly enhanced synergistic increase in these granulocytic subsets, consistent with activation of a feed-forward granulopoietic program. This expansion was significantly impaired in S100A9-deficient cells, where the combinatorial effect of TGF-β and G-CSF was largely abrogated. Instead, S100A9-deficient bone marrow cells preferentially differentiated toward CX3CR1⁺ monocyte/macrophage lineages, indicating a shift in myeloid fate (Figures 3K-3P). Mechanistically, TGF-β robustly induced expression of *S100A9*, *CEBPD*, and *CXCR2*, whereas IL-1β alone was insufficient to drive this program. Notably, co-stimulation with TGF-β and G-CSF further amplified *S100A9* and *CXCR2* expression, reinforcing the cooperative nature of this signaling axis in promoting neutrophil expansion and functional polarization (Figures 3O-3R).

These findings define a TGF-β/S100A9-dependent axis that governs neutrophil expansion and myeloid lineage commitment in metastatic TNBC.

### Metastatic TNBC drives systemic neutrophil expansion and immune remodeling in the spleen

Given the profound differences observed in the tumor microenvironment, we next asked whether metastatic tumors induce systemic immune remodeling. To address this, we performed integrated single-cell RNA sequencing and CyTOF analyses of splenic immune compartments from 4T1, EMT6, and non-tumor-bearing mice. To ensure robust representation of systemic immune states, five independent spleens from EMT6 or 4T1 tumor-bearing mice and four spleens from non-tumor-bearing controls were pooled for single-cell RNA sequencing analyses (Figure S1B).

Unsupervised clustering revealed marked shifts in splenic immune composition in tumor-bearing mice, with 4T1 tumors driving a pronounced expansion of myeloid populations compared to EMT6 and non-tumor-bearing controls (Figures 4A and 4B). Compositional analysis demonstrated a substantial increase in neutrophil populations in the spleens of 4T1 tumor-bearing mice, accompanied by a reduction in lymphoid populations relative to EMT6 and non-tumor-bearing mice (Figure 4C). High-resolution sub clustering identified multiple neutrophil states, including proliferative, immature, inflammatory, and interferon-responsive subsets, with a marked enrichment of immature and inflammatory neutrophil populations in 4T1 spleens (Figure 4D, Table S2A). Notably, 4T1 tumor-bearing mice exhibited a marked enrichment of immature and inflammatory neutrophil populations, consistent with systemic activation of granulopoiesis.

**Figure 4.**
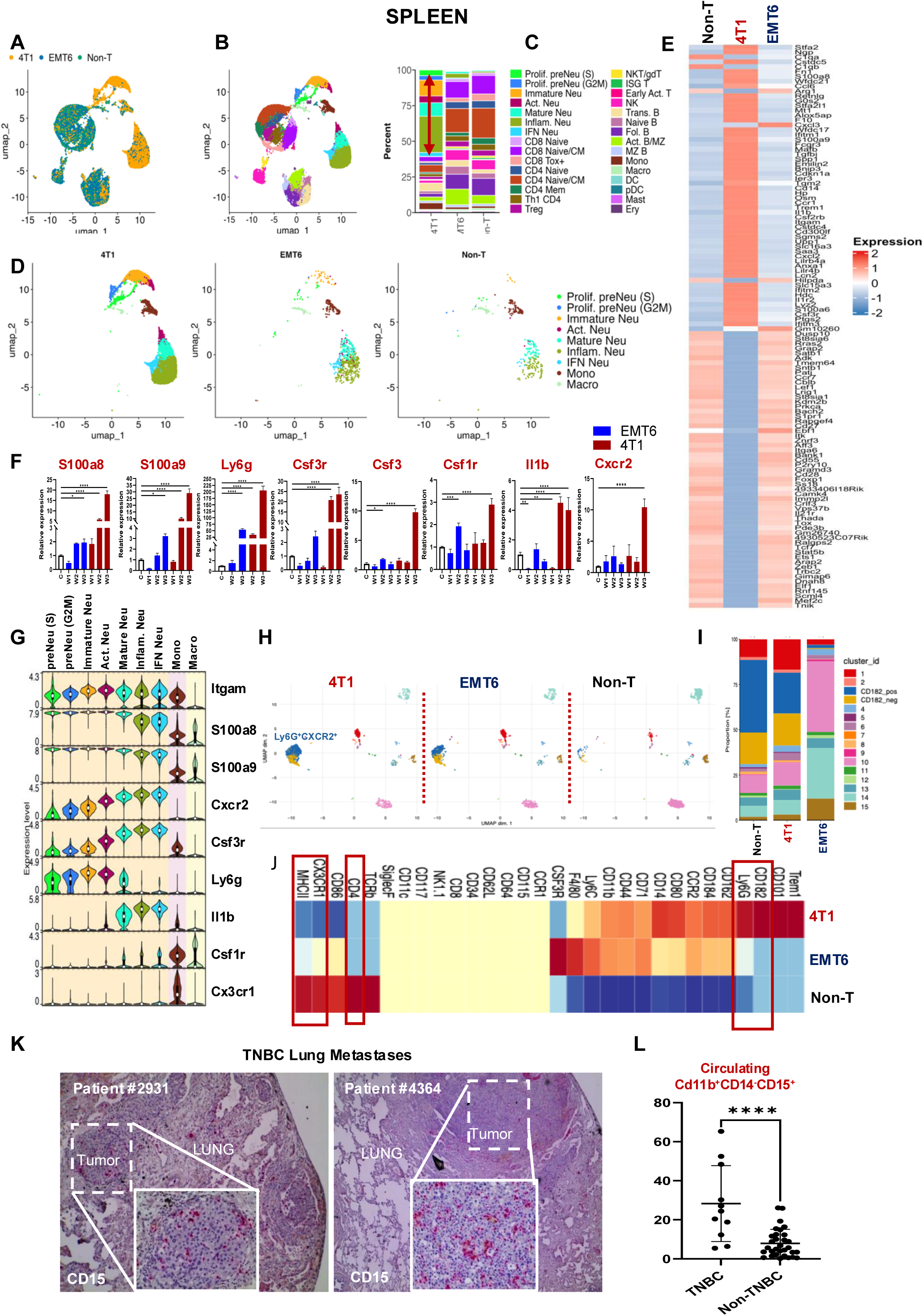
Metastatic TNBC induces systemic neutrophil expansion and splenic immune remodeling. **(A)** UMAP visualization of splenic immune cells from non–tumor-bearing (Non-T), EMT6, and 4T1 tumor-bearing mice, colored by sample origin, demonstrating distinct clustering patterns associated with tumor state. **(B)** UMAP projection colored by annotated immune cell types, identifying major splenic populations including neutrophils, monocytes, macrophages, dendritic cells, and lymphoid subsets. **(C)** Relative proportions of splenic immune populations across conditions, revealing expansion of myeloid compartments, particularly neutrophils, in 4T1 mice compared to EMT6 and Non-T controls. **(D)** Sub clustering of neutrophil and monocyte/macrophage populations, highlighting proliferative pre-neutrophils(preNeu), immature, activated (Act), mature, inflammatory(inflamm), and IFN-responsive neutrophil subsets. **(E)** Heatmap of differentially expressed genes across conditions, showing enrichment of neutrophil-associated and inflammatory transcriptional programs in 4T1 spleens. **(F)** RT-qPCR based quantification of selected neutrophil-associated and inflammatory gene expression (*S100a8, S100a9, Ly6g, Csf3r, Csf3, Csf1r, Il1b, Cxcr2*) across tumors during the tumor growth (n=3, biological replicates) (One-Way ANOVA mean ± SEM test applied compared to Non-T (n=4) *p **<**0.05, ***p **<**0.005, **** p **<** 0.001). **(G)** Violin plots showing expression of key lineage and functional markers (*Itgam, S100a8, S100a9, Cxcr2, Csf3r, Ly6g, Il1b, Csf1r, Cx3cr1*) across splenic immune subsets in EMT6 model. **(H)** UMAP projections of splenic immune cells separated by condition (4T1, EMT6, and Non-T), highlighting expansion of Ly6G⁺CXCR2⁺ neutrophil populations in 4T1 mice. **(I)** Proportional representation of immune cell subsets across conditions. **(J)** Heatmap of CyTOF based immune cell populations, illustrating enrichment of neutrophil-associated signatures in 4T1 and monocyte/macrophage-associated signatures (e.g., *CX3CR1*) in EMT6 and Non-T samples. **(K)** Representative histological images of human TNBC lung metastases showing infiltration of CD15⁺ neutrophils (n=2). **(L)** Quantification of circulating CD11b⁺CD14⁻CD15⁺ granulocytic cells in TNBC (n=11) versus non-TNBC (n=31) patients (Welch-t-test applied for the comparison within conditions **** p **<** 0.001).

Consistent with these observations, differential gene expression analysis revealed increased expression of neutrophil-associated and inflammatory genes, including *S100a8, S100a9, Ly6g, Cxcr2, Csf3r,* and *Il1b*, in 4T1 spleens compared to EMT6 and controls (Figures 4E-4F and Table S2B). In contrast, EMT6 spleens retained higher expression of genes associated with monocyte/macrophage and lymphoid compartments. These transcriptional differences were further supported by lineage-resolved expression patterns across neutrophil subsets (Figure 4G).

Projection of splenic immune subsets across conditions, as defined by CyTOF-based immune profiling, demonstrated preferential accumulation and expansion of neutrophil populations in 4T1 mice, whereas EMT6 and naïve mice exhibited more balanced immune compositions (Figures 4H and 4I). Marker-based heatmap analysis further highlighted enrichment of neutrophil-associated signatures, including CXCR2-driven programs, in 4T1 spleens, while EMT6 spleens maintained higher expression of monocyte/macrophage-associated markers such as CX3CR1 (Figure 4J).

Importantly, these findings were supported by orthogonal validation approaches, where flow cytometry and qPCR analyses were performed on three independent biological replicates per group (Figure S1D), confirming systemic expansion of granulocytic populations.

To extend these findings to human disease, we examined TNBC patient samples and observed enrichment of CD15⁺ myeloid cells in lung metastases (Figure 4K). Consistently, analysis of circulating immune populations revealed a significant increase in CD11b⁺CD14⁻CD15⁺ granulocytic cells in TNBC patients compared to non-TNBC controls (Figure 4L), supporting the clinical relevance of systemic neutrophil expansion.

Together, these findings demonstrate that metastatic TNBC drives systemic immune reprogramming characterized by expansion of immature and inflammatory neutrophil populations, whereas non-invasive tumors preserve a more balanced immune landscape with greater immune activation.

### Integrated trajectory analysis links splenic neutrophil maturation to emergence of CXCR2^hi^ tumor-associated neutrophils in metastatic TNBC

To define the developmental relationship between systemic and tumor-infiltrating neutrophil populations, we performed integrated pseudotime trajectory analysis combining splenic and tumor-derived neutrophils across conditions. UMAP embedding of the combined dataset revealed a continuous developmental continuum spanning proliferative pre-neutrophils, immature and activated states, and terminally differentiated tumor-associated neutrophil (TAN) including CXCR2^hi^ TANs (Figure 5A). Projection of pseudotime across this continuum demonstrated a progressive transition from early proliferative states in the spleen toward terminally differentiated and functionally specialized neutrophil states enriched within tumors, indicating a coordinated systemic-to-local trajectory.

**Figure 5.**
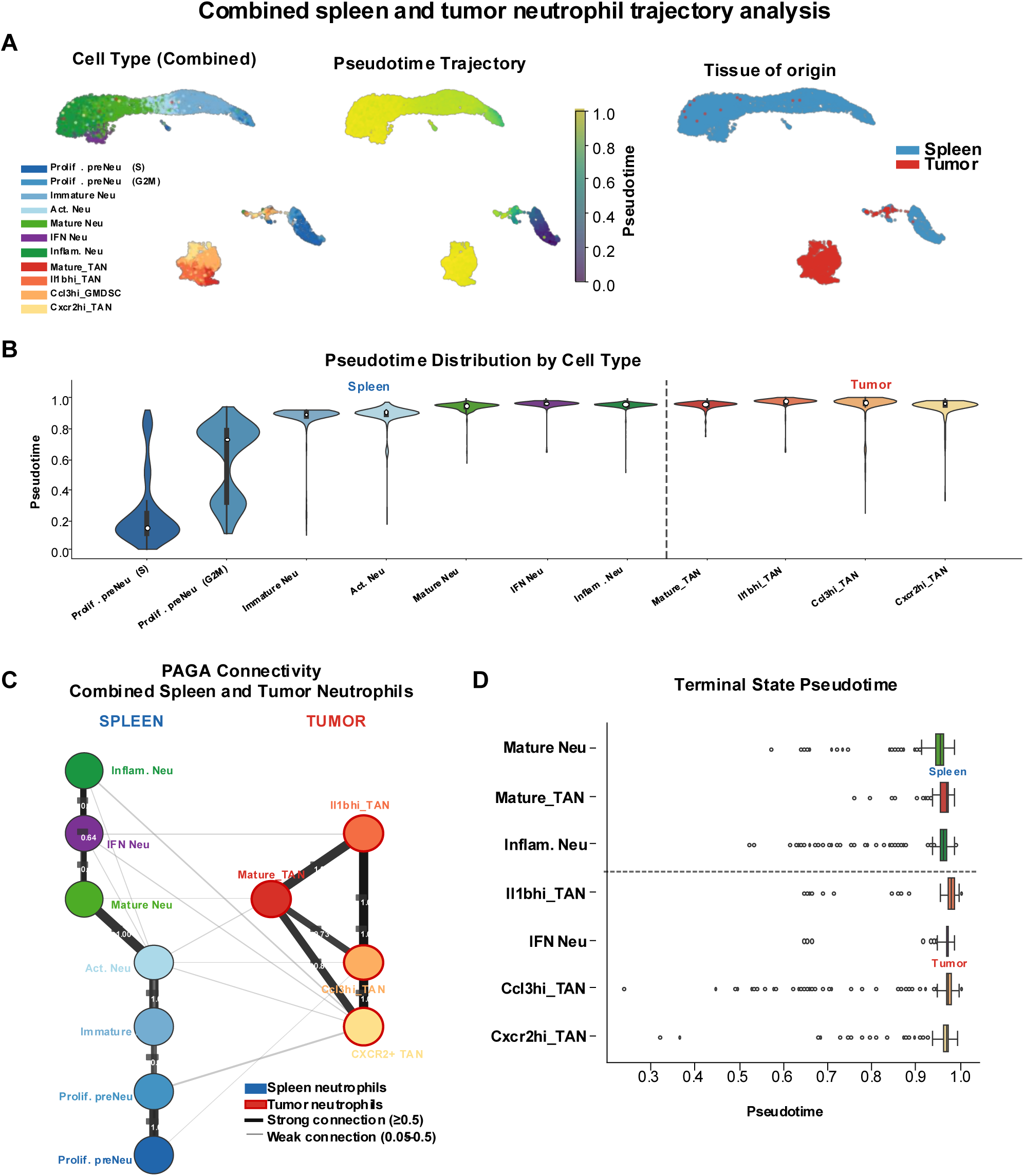
Integrated trajectory analysis links splenic neutrophil maturation to intratumoral reprogramming in metastatic TNBC. **(A)** UMAP embedding of integrated neutrophil populations from spleen and tumor compartments, colored by cell type (left), pseudotime progression (middle), and tissue of origin (right). The trajectory reveals a continuous developmental continuum from proliferative pre-neutrophils (preNeu) and immature neutrophils to terminal tumor-associated states, including mature TANs, Il1b^hi^ TANs, Ccl3^hi^ neutrophils, and Cxcr2^hi^ TAN populations. Spleen-derived cells preferentially occupy early pseudotime states, whereas tumor-derived neutrophils are enriched at late pseudotime. **(B)** Violin plots showing pseudotime distribution across neutrophil states, demonstrating enrichment of early and intermediate states in the spleen and progressive skewing toward terminally differentiated states, including Cxcr2^hi^ TANs, within tumors. **(C)** Partition-based graph abstraction (PAGA) connectivity analysis of combined splenic and tumor neutrophils. Strong directional connectivity links splenic progenitor and intermediate populations to tumor-associated terminal states, with prominent transitions toward mature TANs, Il1b^hi^ TANs, and Cxcr2^hi^ TAN populations, indicating progressive neutrophil reprogramming along a tumor-driven trajectory. **(D)** Terminal state pseudotime analysis demonstrating that tumor-associated neutrophil populations, particularly Il1b^hi^ TANs and Cxcr2^hi^ TANs, occupy the highest pseudotime values, whereas splenic neutrophils remain enriched in earlier developmental states. These data support a model in which systemic neutrophil maturation is coupled to tumor-specific functional polarization.

Quantitative analysis of pseudotime distributions across cell states further highlighted a clear compartmentalization of neutrophil differentiation. Splenic populations were enriched in early and intermediate states, including proliferative preNeu and immature neutrophils, whereas tumor-associated populations were skewed toward late pseudotime states, including mature TANs, Il1b^hi^ TANs, and notably CXCR2^hi^ TANs which occupied the most advanced differentiation states (Figure 5B). Consistent with this, PAGA connectivity analysis revealed a directional trajectory linking splenic neutrophil precursors to tumor-associated terminal states, with strong connectivity between mature TANs, Il1b^hi^ TANs, and CXCR2^hi^ populations, suggesting progressive and structured reprogramming along this axis (Figure 5C). Importantly, CXCR2^hi^ TANs emerged as a terminally differentiated and highly connected node within the tumor compartment, indicating their central role as an endpoint of neutrophil maturation in metastatic tumors.

Terminal state pseudotime analysis further confirmed that tumor-derived neutrophils, particularly CXCR2^hi^ TANs, occupy the highest pseudotime values, whereas splenic neutrophils remain developmentally earlier, consistent with a model of peripheral expansion followed by tumor-specific functional polarization (Figure 5D). This pattern suggests that metastatic tumors do not simply recruit mature neutrophils but instead actively reshape their differentiation trajectory to generate specialized tumor-promoting states.

To further define transcriptional programs underlying this trajectory, we analyzed gene expression dynamics along pseudotime. Early splenic neutrophil states were characterized by expression of genes associated with proliferation and homeostatic functions, whereas late pseudotime states demonstrated induction of inflammatory and tumor-associated programs, including *S100a8, S100a9, Cxcl2,* and *Il1b*, alongside chemokine receptors and migratory programs (Figure S5A). Notably, several of these genes were sharply upregulated in late pseudotime states corresponding to CXCR2^hi^ TANs, reinforcing their identity as tumor-educated, pro-inflammatory neutrophils.

Differential expression analysis of terminal neutrophil subsets further demonstrated that inflammatory and IFN-responsive neutrophils in 4T1 tumors preferentially express genes linked to immunosuppression and pro-tumorigenic signaling, whereas corresponding populations in EMT6 tumors retain features associated with antigen presentation and immune activation (Figure S5B). These findings indicate that tumor context not only drives neutrophil expansion but also dictates their terminal functional states.

Collectively, these data establish a developmental framework in which metastatic tumors co-opt systemic neutrophil ontogeny by promoting expansion of early progenitors and enforcing a maturation trajectory that culminates in CXCR2^hi^ tumor-associated neutrophils, a terminally differentiated population associated with inflammatory signaling, immune suppression, and tumor progression. This coordinated reprogramming links splenic granulopoiesis to intratumoral neutrophil heterogeneity and provides a mechanistic basis for how metastatic TNBC reshapes systemic immunity to support disease progression.

### Tumor-driven expansion of CXCR2⁺Ly6G⁺ neutrophils enforce systemic immune suppression and impair tumor rejection

Building on our observation that metastatic tumors drive neutrophil-biased immune remodeling, we next examined the systemic distribution and functional consequences of these populations across tissues. Flow cytometric analysis revealed a marked expansion of CXCR2⁺Ly6G⁺ neutrophils in 4T1 tumor-bearing mice, evident across tumor, bone marrow, spleen, and lung compartments, with a pronounced increase at 3 weeks post-implantation (Figures 6A-6H). This systemic neutrophil expansion coincided with elevated circulating G-CSF levels (Figure 3G) and increased tumor-derived CXCL2, consistent with a feed-forward granulopoietic axis driving mobilization and peripheral recruitment of CXCR2⁺ neutrophils (Figure 6I).

**Figure 6.**
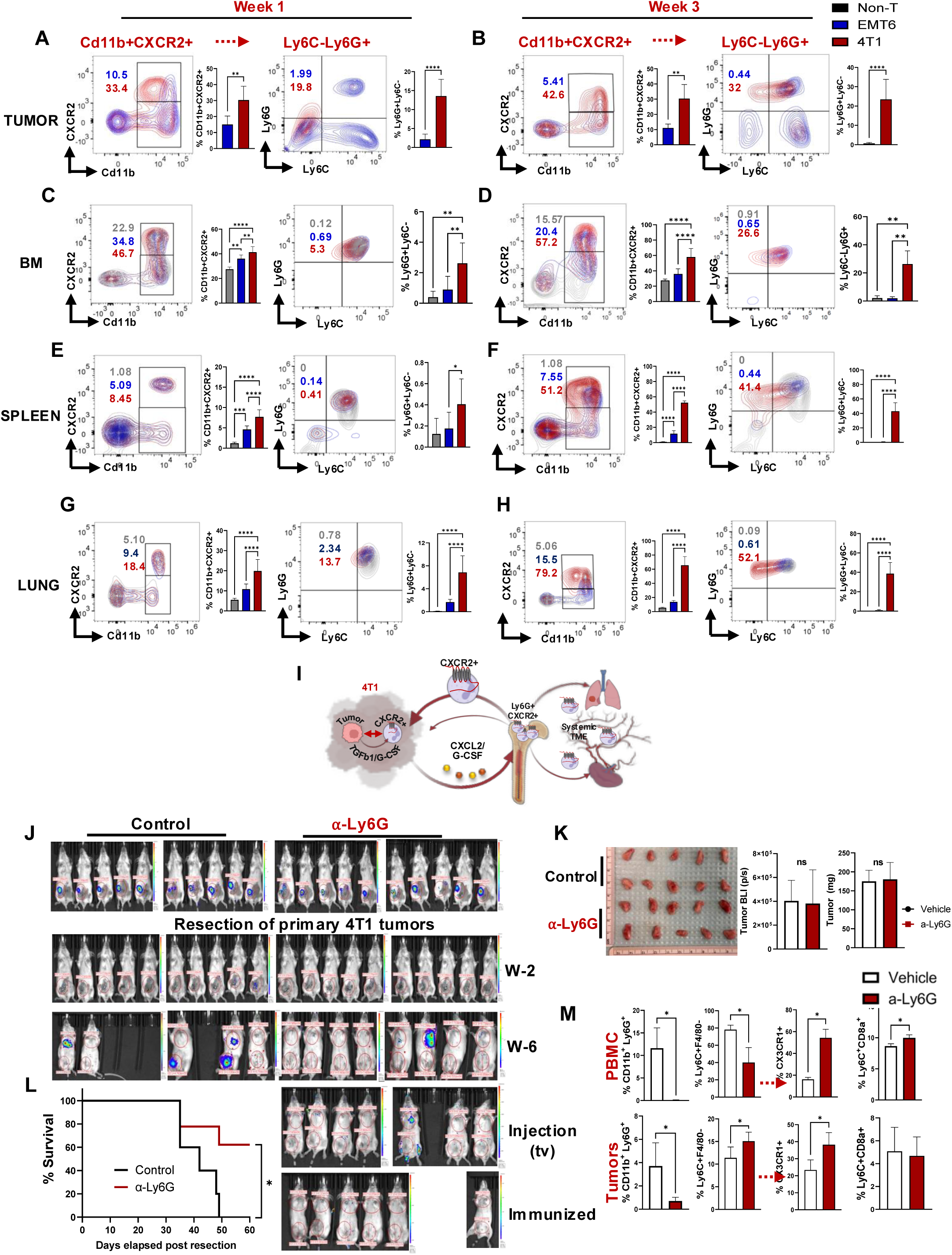
Systemic expansion of CXCR2⁺Ly6G⁺ neutrophils suppress anti-tumor immunity and limits immune memory. (**A, B)** Flow cytometric analysis of tumor-infiltrating CD11b⁺CXCR2⁺ and Ly6G⁺Ly6C⁻ neutrophils at week 1 (n=7, each) and week 3 (n=10, each) in EMT6 and 4T1 tumor models, demonstrating progressive expansion of granulocytic populations in 4T1 tumors (Welch-t-test applied to compare tumor-derived values **p **<**0.01, **** p **<** 0.001). **(C, D)** Bone marrow analysis showing expansion of CD11b⁺CXCR2⁺ progenitors and Ly6G⁺ neutrophils in 4T1 tumor–bearing mice (W1 n=7, W3 n=10) compared to EMT6 (W1 n=7, W3 n=10) and Non-T (n=4) mice (One-Way ANOVA mean ± SEM test applied compared to Non-T values **p **<**0.01, ***p **<**0.005, **** p **<** 0.001). **(E, F)** Splenic immune profiling demonstrating systemic accumulation of CXCR2⁺Ly6G⁺ neutrophils in 4T1 tumor–bearing mice (W1 n=7, W3 n=10) compared to EMT6 (W1 n=7, W3 n=10) and Non-T (n=4) mice (One-Way ANOVA mean ± SEM test applied compared to Non-T values **p **<**0.01, ***p **<**0.005, **** p **<** 0.001).. **(G, H)** Lung analysis showing enrichment of CXCR2⁺Ly6G⁺ neutrophils in pre-metastatic niches of 4T1 tumor–bearing mice (W1 n=7, W3 n=10) compared to EMT6 (W1 n=7, W3 n=10) and Non-T (n=4) mice (One-Way ANOVA mean ± SEM test applied compared to Non-T values **p **<**0.01, ***p **<**0.005, **** p **<** 0.001). **(I)** Schematic model illustrating tumor-derived G-CSF and CXCL2 signaling driving bone marrow mobilization, systemic expansion, and tissue recruitment of CXCR2⁺Ly6G⁺ neutrophils. **(J)** *In vivo* bioluminescence (BLI) imaging of 4T1 tumor–bearing mice treated with control (IgG n=10) or anti-Ly6G (n=10) antibody, before and after resection of primary tumors and following intravenous tumor re-challenge. **(K)** Ex vivo quantification of primary tumor burden showing minimal effect of Ly6G⁺ cell depletion on primary tumor growth. **(L)** KM survival analysis demonstrating improved survival and tumor rejection in neutrophil-depleted mice following tumor re-challenge, indicative of enhanced immune memory (anti-Ly6G treated mice end point 6/10). **(M)** Flow cytometric analysis of PBMCs and tumors showing effective depletion of CD11b⁺Ly6G⁺ neutrophil populations and associated changes in immune composition following anti-Ly6G treatment (n=3 each group) (Welch-t-test applied to compare groups *p **<**0.05).

In contrast, EMT6 tumor-bearing mice failed to induce this granulocytic expansion and instead maintained or expanded monocyte/macrophage populations, characterized by CD115 (CSF1R) expression, across systemic compartments (Figure S4A). These findings highlight fundamentally distinct myeloid programs induced by metastatic versus non-invasive tumors.

To determine the functional relevance of neutrophil expansion, we depleted Ly6G⁺ cells in 4T1 tumor-bearing mice. Anti-Ly6G treatment effectively abrogated systemic accumulation of CXCR2⁺Ly6G⁺ neutrophils (Figures 6J and 6M), without significantly impacting primary tumor burden (Figure 6K). However, following surgical resection of primary tumors, neutrophil-depleted mice exhibited enhanced immune control, with ∼60% of mice rejecting subsequent intravenous 4T1 tumor challenge, indicative of durable anti-tumor immune memory (Figures 6J-6L). In contrast, control-treated mice developed progressive metastatic disease. These findings demonstrate that tumor-driven neutrophil expansion suppresses effective anti-tumor immunity and limits the development of immunological memory.

Given the enrichment of monocyte/macrophage populations in EMT6 tumors, we next assessed their functional role using anti-CD115-mediated depletion. Targeting CD115⁺ cells modestly reduced primary tumor burden in EMT6 models but did not significantly alter systemic immune composition or the establishment of anti-tumor immunity following tumor resection (Figure S4B-S4D). Consistent with our prior findings^11^, EMT6-bearing mice inherently develop protective immunity upon resection of primary tumor, suggesting that monocyte/macrophage populations in this context do not impose systemic immune suppression.

Together, these results demonstrate that metastatic tumors uniquely drive a CXCR2⁺Ly6G⁺ neutrophil-dependent systemic immunosuppressive axis, whereas non-invasive tumors maintain a monocyte/macrophage-dominated immune environment that is permissive to immune activation and memory formation.

### S100A9/Calprotectin signaling drives tumor progression and immune suppression and modulates response to immunotherapy

Having established that metastatic tumors induce a S100A9-dependent neutrophil program that drives systemic immune remodeling, we next investigated the functional consequences of this pathway in tumor progression and therapeutic response. We utilized the E0771 TNBC model, which exhibits higher baseline PD-L1 expression compared to other murine breast cancer models, enabling evaluation of immune checkpoint blockade in the context of S100A9-dependent immune regulation (Figure S5A). In wild-type hosts, anti-PD-L1 treatment resulted in a modest reduction in tumor growth and metastatic dissemination, as assessed by longitudinal bioluminescence imaging (Figures S5B-S5E), indicating partial responsiveness to checkpoint blockade. However, this therapeutic effect was not accompanied by robust systemic immune reprogramming. Flow cytometric analysis of tumor-infiltrating immune cells revealed only limited changes in CD11b⁺CXCR2⁺ neutrophils and modest reductions in Ly6G⁺ granulocytic populations (Figure S5F). Similarly, splenic immune profiling demonstrated selective but incomplete remodeling, with a reduction in CD11b⁺CXCR2⁺ myeloid cells but no consistent expansion of CX3CR1⁺ monocyte/macrophage populations (Figure S5G). These findings suggest that PD-L1 blockade alone is insufficient to fully reverse the neutrophil-dominated systemic immune state.

Strikingly, disruption of S100A9 signaling markedly enhanced responsiveness to immune checkpoint blockade. In S100A9-deficient settings, anti-PD-L1 treatment resulted in significantly reduced tumor burden and metastatic dissemination compared to controls (Figures 7A and 7B), indicating that S100A9-dependent myeloid programming represents a key barrier to effective immunotherapy.

**Figure 7.**
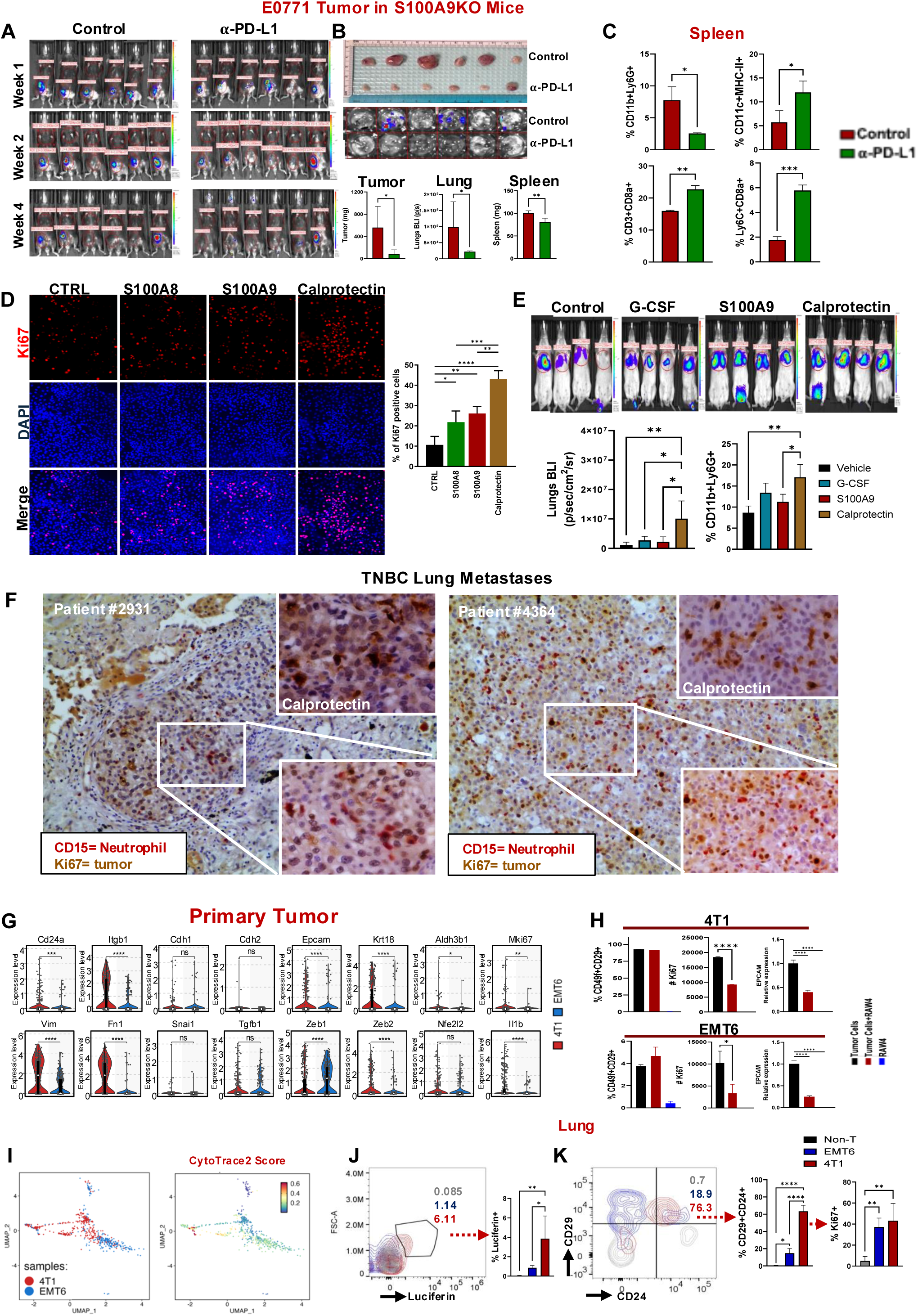
S100A9/calprotectin signaling promotes tumor proliferation, metastatic colonization, and limits response to immune checkpoint blockade. **(A)** *<u>In vivo</u>* BLI of E0771 tumors implanted in S100A9-deficient mice treated with IgG control or anti–PD-L1 antibody, demonstrating enhanced therapeutic response in the absence of S100A9 (n=6 each group). **(B)** Representative images and quantification of tumor burden in primary tumors, lungs, and spleen, showing reduced metastatic dissemination and tumor growth following anti–PD-L1 treatment in S100A9-deficient settings (Welch-t-test applied to compare groups **p **<**0.05, **p **<**0.01). **(C)** Flow cytometric analysis of splenic immune populations, demonstrating reduced neutrophil accumulation and increased antigen-presenting and CD8⁺ T cell responses following PD-L1 blockade (n=3 each group). **(D)** Immunofluorescence analysis demonstrating increased tumor cell proliferation (Ki67) in response to S100A8, S100A9, or calprotectin treatment (experiment was triplicated). Quantification shown at right (Welch-t-test applied to compare groups *p **<**0.05, **p **<**0.01, ***p **<**0.005, **** p **<** 0.001). **(E)** *In vivo* functional assays showing that G-CSF, S100A9, or calprotectin enhances tumor growth and systemic expansion of CD11b⁺Ly6G⁺ neutrophils (n=4, each group) (One-Way ANOVA mean ± SEM test applied compared to calprotectin treated group *p **<**0.05, **p **<**0.01, ***p **<**0.005, **** p **<** 0.001). **(F)** Immunohistochemical analysis of human TNBC lung metastases showing colocalization of CD15⁺ neutrophils with Ki67⁺ proliferating tumor cells and strong calprotectin (S100A8/A9) staining within metastatic lesions. **(G)** Gene expression analysis of primary tumors showing enrichment of epithelial–mesenchymal transition (EMT) and cancer stem cell (CSC)-associated markers in 4T1 tumors compared to EMT6. **(H)** Flow cytometric analysis of tumor cells in the presence and absence of macrophage cell line, Raw264.7, showing increased frequency of CSC-like CD24⁺CD29⁺ populations and enhanced proliferative capacity in 4T1 cells, while EMT6 cells has less CSC-like phenotype with relatively lower proliferation (CSC and Ki67 positivity measued by flow cytometry, EpCAM measued by RT-PCR One-Way ANOVA mean ± SEM test applied compared to tumor cells *p **<**0.05, **** p **<** 0.001) (duplicate experiments were performed twice) **(J)** UMAP projection and CytoTRACE analysis demonstrating increased stemness and differentiation potential in 4T1 tumor cells relative to EMT6. **(J-K)** Flow cytometric quantification of luciferase⁺ tumor cells, indicating increased abundance of tumor cells with high proliferation capacity (Ki67+) in 4T1-bearing mice (n=5) compared to EMT6 (n=5) and Non-T (n=4) mice at week 3 of tumor growth (One-Way ANOVA mean ± SEM test applied compared to Non-T values **p **<**0.01, ***p **<**0.005, **** p **<** 0.001).

To further define the functional role of S100A8/A9 (calprotectin), we examined its direct impact on tumor cell behavior. Exposure to recombinant S100A8, S100A9, or calprotectin significantly increased tumor cell proliferation, as measured by Ki67 staining, with calprotectin exerting the most pronounced effect (Figure 7D). In vivo, administration of G-CSF, S100A9, or calprotectin enhanced pulmonary tumor growth and promoted systemic expansion of granulocytic populations, supporting a feed-forward loop linking tumor-derived inflammatory cues to neutrophil mobilization and tumor progression (Figure 7E).

Consistent with these findings, analysis of human TNBC lung metastases revealed abundant infiltration of CD15⁺ neutrophils in close proximity to Ki67⁺ proliferating tumor cells, accompanied by strong calprotectin expression, highlighting the clinical relevance of this axis in metastatic disease (Figure 7F).

At the primary tumor level, transcriptional profiling demonstrated that metastatic 4T1 tumors exhibit increased expression of epithelial and stemness-associated markers, including *Epcam*, *Krt18*, *Aldh1a1*, and *Mki67*, alongside induction of mesenchymal and inflammatory programs compared to EMT6 tumors (Figure 7G). Co-culture of tumor cells with monocyte/macrophage populations suppressed tumor cell proliferation, suggesting that non-neutrophilic myeloid compartments may exert context-dependent anti-tumor effects (Figure 7H).

Single-cell-based CytoTRACE2 analysis further demonstrated enrichment of less differentiated, stem-like tumor cell states in 4T1 tumors relative to EMT6 (Figure 7I). Functionally, these differences translated into increased tumor-initiating capacity, as luciferase-labeled 4T1 cells exhibited enhanced metastatic colonization and growth in the lung (Figure 7J). Flow cytometric analysis confirmed increased tumor burden and expansion of proliferative cancer stem-like cells (CD24⁺CD29⁺) in 4T1-derived lung metastases compared to EMT6 (Figure 7K).

Collectively, these findings demonstrate that S100A9/calprotectin signaling promotes tumor progression through dual mechanisms: (i) establishment of a neutrophil-dominated, systemically immunosuppressive microenvironment that limits effective anti-tumor immunity, and (ii) direct enhancement of tumor cell proliferation, stemness, and metastatic fitness. Importantly, targeting this axis enhances responsiveness to immune checkpoint blockade, identifying S100A9-dependent myeloid programming as a therapeutically actionable pathway in metastatic TNBC.

## Discussion

Our study establishes a systems-level framework linking tumor-intrinsic state to coordinated local and systemic immune remodeling^16,17^ in TNBC through a direct comparative analysis of metastatic versus non-invasive tumor models. By integrating single-cell profiling, functional perturbation, and clinical datasets, we identify a metastasis-specific TGF-β/S100A9/CXCR2 axis that drives neutrophil expansion, immune suppression, and cancer stem cell (CSC) plasticity in aggressive disease, while revealing a distinct immune trajectory in non-invasive settings characterized by preserved monocyte/macrophage and T cell compartments. Central to this approach is the use of EMT6 as a non-invasive counterpart to the highly metastatic 4T1 model. EMT6 was specifically selected for its immunologically “hot” phenotype and its capacity to sustain systemic anti-tumor immunity^11^, enabling a biologically informative contrast with the immunosuppressive and metastasis-prone 4T1 tumors. While non-metastatic variants such as 67NR share a common genetic background with 4T1, EMT6 provides a complementary system with a distinct immune composition, particularly enriched for CX3CR1⁺ monocyte/macrophage populations and functional T cell responses, allowing us to more clearly delineate tumor-driven differences in immune programming. Together, this comparative framework reveals that immune remodeling is not a uniform consequence of tumor growth but is instead specified early by tumor state, dictating divergent systemic trajectories that ultimately govern metastatic potential and therapeutic responsiveness.

Mechanistically, our data identify a TGF-β/C/EBPδ/S100A9 regulatory axis as a key driver of neutrophil reprogramming. TGF-β robustly induced expression of *S100A9*, *Cebpd*, and *Cxcr2*, consistent with prior work demonstrating that C/EBPδ governs epigenetic activation of the *S100a8* and *S100a9* loci during inflammatory myelopoiesis^18^. While our data highlight induction of *S100A9*, it remains possible that coordinated upregulation of *S100A8* also contributes to the observed phenotypes through formation of the S100A8/A9 heterodimer^19^. In either scenario, these findings reinforce a dominant role for calprotectin-mediated signaling in linking TGF-β-driven myelopoiesis^20,21^ to neutrophil functional polarization. Importantly, our findings are mechanistically distinct from previous studies showing that TGF-β promotes polarization of tumor-associated neutrophils toward a pro-tumorigenic phenotype, primarily through immune suppressive effector functions and inhibition of cytotoxic T cell activity. In contrast, our study identifies an upstream TGF-β-driven developmental program that actively rewires systemic granulopoiesis through induction of the C/EBPδ/S100A9 axis, resulting in expansion of CXCR2^hi^ immature neutrophils and coordinated local and systemic immune remodeling.

A key strength of this study lies in the comparative analysis of metastatic and non-invasive TNBC models, which provides a unique framework to disentangle tumor-intrinsic versus immune-driven determinants of disease progression. While most high-dimensional studies from pre-clinical and clinical settings have focused exclusively on advanced late-stage tumors^22–24^, our findings demonstrate that immune remodeling is initiated through early tumor-immune crosstalk, whereby tumor-intrinsic programs actively instruct immune cell fate and function from the onset of tumor development. Importantly, this early crosstalk extends beyond the local tumor microenvironment to shape systemic immune architecture, as metastatic tumors rapidly engage bone marrow and splenic compartments to drive granulopoiesis and neutrophil mobilization.

In contrast to the neutrophil-dominated, immunosuppressive landscape observed in metastatic 4T1 tumors, the EMT6 model retained a myeloid compartment enriched for CX3CR1⁺ monocyte/macrophage populations, suggesting a fundamentally distinct immune trajectory associated with tumor restraint. CX3CR1 has emerged as a key marker of differentiated, patrolling, and tissue-resident monocyte/macrophage subsets that support immune surveillance and antigen presentation, and its expression has been associated with favorable immune contexts across multiple malignancies. In particular, CX3CR1⁺ immune populations have been linked to enhanced cytotoxic T cell priming and improved responsiveness to immunotherapy. Notably, recent studies have identified CX3CR1-positive immune populations as key mediators of effective T-cell responses and durable tumor control, with CX3CR1 expression serving as a marker of productive anti-tumor immune activation and favorable clinical outcome^25,26^. Our TCGA analyses further support these observations, demonstrating that CX3CR1 expression is reduced in TNBC compared with ER-positive breast cancers and progressively declines with increasing tumor grade. Furthermore, CX3CR1 expression on CD8⁺ T cells have been identified as a robust biomarker of effective anti-tumor immunity and clinical response to immune checkpoint blockade, reflecting a differentiated effector state enriched for tumor-reactive clones^27^. Notably, expansion of circulating CX3CR1⁺ CD8⁺ T cells correlate with response and survival in non-small cell lung cancer patients receiving anti-PD-1 therapy^28^, highlighting CX3CR1 as a dynamic indicator of anti-tumor immune engagement.

Importantly, our findings highlight that these differences emerge at early stages of tumor development and extend beyond the local tumor microenvironment to shape systemic immune architecture. Metastatic tumors rapidly engage bone marrow and splenic compartments to drive emergency granulopoiesis and neutrophil mobilization, whereas non-invasive tumors fail to induce such systemic reprogramming. This early divergence in both local and systemic immune states likely represent a critical determinant of metastatic progression and therapeutic response, underscoring the importance of studying tumor-immune interactions across multiple compartments and disease stages.

While prior studies have characterized the immunosuppressive tumor microenvironment in TNBC^3,4^, these analyses have largely focused on established metastatic disease. Our comparative approach demonstrates that immune remodeling is not uniform but instead dictated by tumor state. Metastatic 4T1 tumors induce a systemic, neutrophil-dominated program, whereas non-invasive EMT6 tumors preserve immune activation and memory. These findings extend previous work showing that tumors can reprogram hematopoiesis^9^ and elevate neutrophil-to-lymphocyte ratios associated with poor prognosis^10^, by linking these systemic effects directly to metastatic competence.

We identify early activation of TGF-β and G-CSF signaling as key drivers of emergency granulopoiesis, leading to expansion of immature CXCR2⁺ neutrophils. This is consistent with prior studies showing G-CSF-mediated mobilization of suppressive neutrophils^9,29^, but we further demonstrate that this process is mechanistically dependent on S100A9. Importantly, pseudotime analysis reveals that metastatic tumors do not generate new differentiation programs but instead hijack a conserved neutrophil developmental trajectory, skewing it toward immature and inflammatory states. This aligns with emerging concepts that tumor-induced myelopoiesis produces functionally distinct neutrophil subsets with immunosuppressive properties^5,7^.

Our data identify S100A9 as a critical node integrating tumor-derived signals with systemic immune remodeling. Previously S100A9 has been shown to inhibit dendritic cell differentiation and promote expansion of myeloid-derived suppressor cells (MDSCs) and macrophages in cancer^15,30,31^, supporting our findings that calprotectin-dependent signaling drives systemic myeloid remodeling and immunosuppression in metastatic TNBC. It was also reported that S100A8/A9 (calprotectin) has been implicated in MDSC function through TLR4 and RAGE signaling^15,32^. However, it should be noted that the biological activity of S100A8 and S100A9 is highly dependent on their oligomerization state^33^. While S100A8/A9 heterodimers can engage receptors such as TLR4, higher-order tetramer formation has been reported to attenuate this activity^33^. Therefore, the functional effects observed in our assays may reflect contributions from both heterodimeric and homodimeric S100 species depending on protein source and biochemical composition, an important consideration when interpreting calprotectin-mediated signaling. Our findings extend this role by demonstrating its necessity for CXCR2⁺ neutrophil expansion and functional polarization.

Loss of S100A9 not only impairs CXCR2^+^ neutrophil expansion but also redirects myeloid differentiation toward CX3CR1⁺ monocyte/macrophage populations, highlighting its role in lineage commitment. This dual function positions S100A9 as both a regulator of myelopoiesis and an effector of immune suppression.

It is important to note that S100A9-deficient mice are functionally deficient in both S100A9 and S100A8 at the protein level due to the obligate stabilization of the S100A8/A9 heterodimer, despite preserved S100A8 mRNA expression. Thus, the phenotypes observed here likely reflect disruption of the S100A8/A9 (calprotectin) axis rather than S100A9 alone.

A key conceptual advance of this study is the integration of immune remodeling with tumor cell plasticity. We show that neutrophil-derived S100A8/A9 promote EMT and MET-associated CSC states, processes previously linked to inflammatory signaling^8^.

This supports a model in which neutrophils act not only as suppressors of adaptive immunity but also as active regulators of tumor evolution, facilitating both dissemination and colonization. The observation that calprotectin directly enhances tumor cell proliferation further underscores the functional coupling between immune and tumor compartments.

While immune checkpoint inhibitors have shown limited efficacy in TNBC^34,35^, our data demonstrate that S100A9 ablation significantly enhances response to anti-PD-L1 therapy. Previously S100A9 has been shown to inhibit dendritic cell differentiation and promote expansion of myeloid-derived suppressor cells (MDSCs) in cancer^31^, supporting our findings that calprotectin-dependent signaling drives systemic myeloid remodeling and immunosuppression in metastatic TNBC. This suggests that neutrophil-driven immune suppression^36^ represents a key barrier to immunotherapy. Targeting components of this axis, including CXCR2, G-CSF signaling, or S100A9, may therefore represent a rational strategy to sensitize tumors to checkpoint blockade. Indeed, CXCR2 inhibitors are currently under clinical investigation and may be particularly effective in neutrophil-high tumors.

Our findings align with emerging evidence that tumor-infiltrating neutrophils converge on a limited number of terminal states regardless of their bone-marrow origin. Recent pan-cancer single-cell atlases revealed that neutrophils across 17 human tumor types segregate into a conserved set of subsets, including an HLA-DR^+^ antigen-presenting population whose abundance tracks with favorable prognosis^37^, while in vivo lineage tracing showed that both immature and mature neutrophils are deterministically reprogrammed within hypoxic tumor regions toward a shared pro-angiogenic terminal state^38^. A high-resolution NSCLC atlas similarly identified tissue-resident neutrophil subpopulations whose transcriptional signatures predict failure of anti-PD-L1 therapy^39^. Our pseudotime analyses extend these observations to TNBC, demonstrating that splenic neutrophil precursors mobilized by metastatic 4T1 tumors feed an analogous terminal CXCR2^+^CXCL2^+^S100A9^+^ tumor-associated neutrophil state. This convergence argues against targeting differentiated neutrophils alone and reinforces the rationale for therapeutically engaging upstream regulators, TGF-β, C/EBPδ, and S100A, that specify the immunosuppressive program.

A recent companion study reported that mammary tumors reprogram the hematopoietic stem and progenitor cell compartment via IL-1β and nucleotide-derived signaling to bias bone-marrow output toward immunosuppressive granulopoiesis, and that IL-1β blockade reverses this bias and reduces metastatic spread^40^. Our identification of a TGF-β/C/EBPδ/S100A9 axis acting early in the bone-marrow compartment of 4T1-bearing mice complements that work and nominates an additional, IL-1β-independent module driving emergency granulopoiesis. Together, these findings define a converging logic in which distinct tumor-derived cues funnel into transcriptional and epigenetic rewiring of HSPCs to license metastasis-supporting myeloid output, and they argue that combinatorial targeting of these upstream nodes may yield broader and more durable effects than blockade of any single pathway.

From a translational standpoint, our data reinforce S100A8/A9 as a clinically actionable axis in TNBC. S100A8/A9 expression has been shown to stratify early-stage TNBC patients by recurrence risk and to identify tumors that respond to combined PIM kinase and PD-1/PD-L1 inhibition in preclinical models^41^, providing a tractable pharmacological entry point that aligns with our genetic S100A9 ablation data. In parallel, physical interaction of neutrophils with tumor cells through CX3CR1^+^ macrophage-orchestrated niches has been shown to directly enhance breast-cancer aggressiveness and to predict worse survival^42^, consistent with the calprotectin-driven CSC-plasticity loop we describe. Taken together, these convergent lines of evidence position the TGF-β/C/EBPδ/S100A9 axis as a therapeutically tractable node whose pharmacological inhibition, alone or in combination with immune-checkpoint blockade, merits prioritized clinical evaluation in neutrophil-high TNBC subsets.

In contrast to the dominant role of neutrophils in metastatic tumors, EMT6 tumors rely on monocyte/macrophage populations that do not suppress systemic immunity and may support anti-tumor responses. This highlights the functional heterogeneity of myeloid cells and underscores the need for context-specific therapeutic strategies. Our findings are consistent with prior work demonstrating distinct roles for monocytic versus granulocytic MDSCs in tumor progression^12^, but extend this concept by linking these populations to tumor state and systemic immune regulation.

Collectively, our findings support a model in which metastatic tumors actively reprogram systemic myelopoiesis through a TGF-β/C/EBPδ/S100A8/A9 axis, driving expansion and mobilization of CXCR2⁺ neutrophils that undergo further functional polarization within the tumor microenvironment. This coordinated local and systemic remodeling establishes an immunosuppressive niche that promotes cancer stem cell plasticity and metastatic progression. In contrast, non-invasive tumors fail to engage this axis and instead maintain CX3CR1⁺ monocyte/macrophage populations that support immune surveillance. Targeting this neutrophil-centric axis therefore represents a tractable strategy to uncouple immune suppression from metastatic progression.

## Acknowledgements

We acknowledge the Microscopy, Imaging and Cytometry Resources Core (MICR) at Wayne State University for the flow cytometry support. MICR is supported in part by the NIH Center grant P30 CA22453 to the Karmanos Cancer Institute. This study was supported by the National Institute of Health NCI grant R01CA251676 and Karmanos Cancer Institute Startup fund to H. Korkaya. NIH 1R01CA282098-01A1 and the Breast Cancer Research Foundation Funds, BCRF –24-173 to MW. NIH R01CA264983 to HS, and Evans County CARES, Inc. to H.K. and H.S.

## Author Contributions

Conceptualization, F.K.A., M.S.W., H.S., and H.K.; Methodology, F.K.A., A.B.C., H.K.A., C.J., E.L., R.P., M.A., E.B., A.A., A.L.G., T.V., G.D., A.C., M.G., S.K.-B., H.A., N.N., G.O., R.B., and C.C.H.; Investigation, F.K.A., A.B.C., H.K.A., C.J., E.L., R.P., M.A., E.B., A.A., and A.L.G.; Formal Analysis, F.K.A., A.B.C., G.D., and M.A.; Resources, T.V., S.K.-B., R.B., M.S.W., H.S., and H.K.; Writing, Original Draft, H.K.; Writing, Review & Editing, all authors; Supervision, M.S.W., H.S., and H.K.; Funding Acquisition, M.S.W., H.S., and H.K.

## Inclusion and Diversity

We support inclusive, diverse, and equitable conduct of research. One or more of the authors of this paper self-identifies as a member of an underrepresented ethnic minority in their field of research or within their geographical location.

## Data and Code Availability

Single-cell RNA-sequencing and CyTOF data generated in this study are being deposited in the Gene Expression Omnibus and FlowRepository, with accession numbers to be added at acceptance. Original code is available on GitHub and will be activated upon publication. Any additional information required to reanalyze the data reported in this work is available from the lead contact upon reasonable request.

## RESOURCE AVAILABILITY

### Lead contact

Request about this study and for resources should be directed to the lead contact, Hasan Korkaya

### Materials availability

No new materials were generated for this study. S100A0KO mice in the key sources table were previously acquired by H.K. from T.V. via a material transfer agreement and can be requested by the lead contact.

### Data and code availability

- The TCGA data used in this study are based on data generated by the TCGA Research Network (http://cancergenome.nih.gov/) and are available in a public repository on the cBioportal for Cancer Genomics website (http://www.cbioportal.org/).
- Single-cell RNA Sequencing data from mouse tumors and spleens are available at Gene Expression Omnibus (GEO) and the accession number is listed in the key resource table.
- Custom analysis scripts and codes for figure generation are publicly available. Repositories are listed in the key resources table.
- All other data supporting the study’s findings are available in the article and the Supplemental Documents, and from the lead contact upon request.

## STAR★METHODS

Detailed methods are provided in the online version of this paper and include following:

- KEY RESOURCES TABLE
- EXPERIMENTAL MODEL AND STUDY PERTICIPANT DETAILS
  ○ Human patient samples
  ○ Animals
  ○ Tumor cell lines
  ○ Lentiviral transduction of tumor cell lines
- METHOD DETAILS
  ○ *In vivo* experiments in mouse tumor models, bioluminescent imaging, and treatments
  ○ Tissue dissociation
  ○ *Ex vivo* mouse bone marrow stimulation experiments
  ○ Immune profiling by Flow Cytometry
  ○ RNA isolation and quantitative real-time PCR (RT-qPCR)
  ○ Immunohistochemistry (IHC) and immunofluorescence (IF) staining
  ○ Time-of-flight mass spectrometry (CyTOF)
  ○ Single –cell RNA sequencing (ScRNA-req)
- QUANTIFICATION AND STATISTICAL ANALYSIS
  ○ Time-of-flight mass spectrometry (CyTOF) data processing
  ○ Single-cell RNA sequencing data processing
  ○ Neutrophil trajectory analysis
  ○ Statement about AI tool use
  ○ Statistical analysis

## EXPERIMENTAL MODEL AND STUDY PARTICIPANT DETAILS

### Human specimens

Whole blood samples of patients diagnosed with breast cancer were collected by Augusta University Health Sciences Department with informed consent. All analysis of human peripheral blood mononuclear cells (PBMCs) were performed under the protocol (Title: Improving effectiveness of immunotherapy by targeting the microenvironment in preclinical models of breast cancer) approved by the review board of IRB: **1315482-1**

### Animals

Animal studies were performed in accordance with the Institutional Animal Care and Use Committee (IACUC) Wayne State University. The animal protocol number IACUC-23-05-5789 was reviewed and approved by Division of Laboratory Animal Resources (DLAR). Balb/c and C57BL/6 mice were purchased from the Jackson Laboratory and S100A9-KO mice were kindly provided by Prof. Thomas Vogl^33^. Homozygous S100A9-KO female mice were generated by breeding female and male heterozygous mouse to generate homozygous for S100A9 deletion. Mice were housed at pathogen-free rooms with 50–70% humidity, and 12/12 h light/dark cycle. 6-to-8 weeks old female mice were used in all experiments. The threshold for tumor volume did not exceed 1500mm3 in all animal experiments.

### Tumor cell lines

4T1, EMT6, AT3, E0771 and Raw264.7 cell lines were purchased from ATCC. All cell lines periodically tested for mycoplasma contamination using a PCR-based detection kit (ATCC). 4T1, EMT6, AT3 and E0771 cell lines were transfected with luciferase-expressing lentivirus to generate stable cell lines and monitor tumor growth and metastatic formation in animals. 4T1, 4T1-Luc, EMT6 and EMT6-Luc cell lines were maintained in RPMI media (Gibco) supplemented with 10%FBS (Cytiva) and 1% antibiotic/antimycotic (100X) (Gibco). AT3 and E0771 cell lines were maintained in DMEM supplemented with 10% FBS and 1% antibiotic/antimycotic. Raw264.7.1 cell line was maintained in DMEM supplemented with 10% FBS and 1% penicillin-streptomycin (Gibco). Cells were maintained at 37 ℃ in a humidified 5% CO_2_ atmosphere incubator.

### Lentiviral transduction of tumor cell lines

Cell lines were seeded at the density of 5×10^4^ per well on 6 wells and incubated overnight in cell culture incubator. Cells were transduced with 200,000TU/well firefly luciferase lentivirus (BPS79692-G, BPS Bioscience) in fresh cell culture media for 24 hours. After 24 hours of transduction, the medium was changed to regular growth media.

## METHOD DETAILS

### *In vivo* experiments in mouse tumor models, bioluminescent imaging, and treatments

For orthotopic 4T1-Luc, EMT6-Luc, AT3-Luc and E0771-Luc cells, mice were anesthetized with Ketamine/Xylazine cocktail at the dose of 80-100mg/kg-10-12.5mg/kg/kg. 1×1cm L-shaped incision was made between the abdominal midline and the 4^th^ and 5^th^ mammary fat pads. Skin was opened with sterile cotton swaps to expose the 4th mammary fat pad. 5×10^4^ 4T1-Luc or EMT6-Luc cells on media containing %50 Matrigel were implanted in Balb/c mice. 1×10^5^ AT3-Luc or E0771-Luc cells were implanted in C57BL/6 and/or S100A9-KO mice. 5mg/kg Rimadyl/Carprofen (Zoetis) were administered to mice before and the surgery and 24hr thereafter. For bioluminescent imaging, mice were anesthetized by inhalation of 2% isofluorane in oxygen and injected 150mg/kg D-luciferin (Revvity). Mice were imaged 10 minutes later using IVIS Spectrum (Pelkin Elmer). For Ly6G blockade, 200 μg/ml of anti-mouse Ly6G (BE0075, BioXCell); for PD-L1 blockade, 200 μg/ml anti-mouse PD-L1 (BE0101, BioXCell) antibody; and for CSF1R blockade, 250 μg/ml anti-mouse CD115 (BE0213, BioXCell) were injected intraperitoneally (i.p.) every other day for three weeks. For tail vein injections, 1×10^5^ EMT6-Luc tumor cells were resuspended in 200μL of sterile PBS and injected through the tail vein. Animals were imaged 1 hour after tail vein injection to confirm tumor cells localized in the lungs, then monitored weekly for metastatic progression. For survival analysis, established primary tumors and draining lymph nodes were resected from the 4^th^ mammary fat pat by cauterizing the surrounding arteries. Animals were monitored weekly by residual tumor growth and metastatic formation. In all experiments, control mice were injected with 0.9% Sodium Chloride otherwise indicated. Skin flaps were sutured by metal clips after mammary fat pat surgery. At the end points of each experiment, mice were euthanized by CO_2_ asphyxiation, and tissues were collected for further analysis.

### Tissue dissociation

Single cell suspension was prepared as previously described^43^. Briefly, whole blood was collected into K2/EDTA Vacutainer (BD) tubes by cardiac puncture right after euthanasia. Spleens were excised, weighted, and imaged and then smashed through the 70μm strainers (Fisher Scientific) using the plunger side of the syringe. Tumor and lungs were excised, weighted, lungs were imaged and minced with forceps, and enzymatically digested in DMEM containing 1X Collagenase/Hyaluronidase (Stem Cell Technologies) and 0.25mg/ml DNase-I (Roche) at 37℃ water bath for 45 minutes. Suspensions were filtered through 70μm strainers to retrieve single cells. Red blood cells were eliminated by the ACK Lysing buffer (Gibco) according to manufacturers’ instructions. Single cells were resuspended in D-PBS (Gibco) for further analysis, otherwise stored at –80℃ as a pellet by 300xg 5min 4℃ centrifugation.

### Immune profiling by Flow Cytometry

Single cells from dissociated tissues were counted, and 1.5×10^5^ cells were taken into 96 well plates. Cells were washed twice with D-PBS and incubated with fixable viability dye (Ghost Dye Red 710 1:2000, Fixable viability dye 510/450 1:1000) for 15 minutes at room temperature. Cells were blocked in FcX PLUS (156604, Biolegend) at room temperature for 10 minutes. For extracellular staining, samples were incubated with antibody cocktails in dark for 30 minutes at 4℃ in Stain buffer (BD Biosciences) containing Brilliant Stain Buffer (BD Biosciences) for 3 different optimized flow cytometry panels for lymphocytes, myeloid cells and cancer stem cells. For intracellular staining, cells were fixed with IC Fixation (00-8222-49, Invitrogen) buffer for 15 minutes at room temperature and then washed twice with 1X permeabilization buffer (00-5523-00, Invitrogen). Intracellular antibody staining was performed at room temperature for 30 minutes with antibody cocktails prepared in a 1X permeabilization buffer containing 2% FBS. For nuclear staining, cells were incubated at room temperature for 15 minutes in Fix/Perm buffer (00-5523-00, Invitrogen) and washed twice with 1X permeabilization buffer (00-5523-00, Invitrogen). Nuclear antibody staining was performed at 4℃, overnight in dark with antibody cocktails in 1X permeabilization buffer containing 2% FBS. For cancer stem cell assays, 1.5×10^5^ cells were stained for cell surface antibodies Stain buffer (BD Biosciences) for 30 minutes at 4℃ in dark and incubated with impermeant viability dye, DAPI (BD Bioscience) for 5 minutes on ice. Fluorescent dye conjugated antibodies that were used in the experiments were listed in the key resources table. Samples were run in Northern Lights Full Spectrum Flow Cytometry (Cytek) using the plate-reader at medium speed. Runs were analyzed on FlowJo 10.10.

### *Ex vivo* mouse bone marrow stimulation experiments

Bone marrow cells were isolated from femur and tibia from 8-10 weeks old WT (Balb/c and C57BL/6) and S100A9 knock-out mice. Briefly, mice were euthanized with CO_2_ asphyxiation and sprayed with 70% ethanol. Hind limbs were exposed off the skin and dislocated from the hip joint and placed into ice-cold D-PBS (Gibco). Muscles were removed to expose femur and tibia using forceps and scalpel. Femur and tibia were cut at either end. bone marrows were flushed using a 25G needle with 10mL ice-cold D-PBS under sterile tissue culture hood. Disaggregated bone marrow collected and passed through 70μm strainers (Fisher Scientific) into a sterile 50ml centrifuge tube and kept on ice while samples were counted. Bone marrow cells were seeded with a density of 1×10^6^ on 6-well plates (Corning, 3471). DMEM with 10% FBS without antibiotics were supplemented with G-CSF (100ng/ml), GM-CSF (20ng/ml), TGF-b (5ng/ml), S100A9 (50 ng/ml), S100A8 (50 ng/ml), Carprofen (50 ng/ml), or LPS (100ng/ml) as indicated in the figures. 1ml Fresh media added every two days with supplements. Cells were collected at the day 7 of the differentiation by washing with ice-cold D-PBS and pellets were collected by centrifugation at 300xg for 5min at 4 ℃ for further applications.

### RNA isolation and quantitative real-time PCR (RT-qPCR)

Total RNA was isolated from tissue-derived single cells and in vitro cell lysates using QIAwave RNA Mini Kit (74536, Qiagen). cDNA was generated using iScript Advanced cDNA Synthesis Kit (1725038, Bio-rad). Quantitative RT-PCR was performed in QuantStudio5 Real-time PCR System (Applied Biosystems). iTaq universal SYBR Green SuperMix (1725121, Bio-Rad) was used in 1X concentration reaction was performed at 40 cycles with 50ng of template cDNA following the steps of: Step1 for 30 sec at 95℃, step 2 for 40 times of 15 sec at 95℃ and 30sec at 60℃, step 3 for melt curve for 1 sec at 95℃, 20 sec at 65℃ and 1 sec at 95℃. The primer sequences for qRT-PCR were listed in the key resources table 2. Actin was used as a housekeeping gene, and no template controls were included in all experiments. All samples were run in triplicate in two biological replicates. The relative mRNA expression was calculated using 2-ΔΔCT method.

**Table 1:**
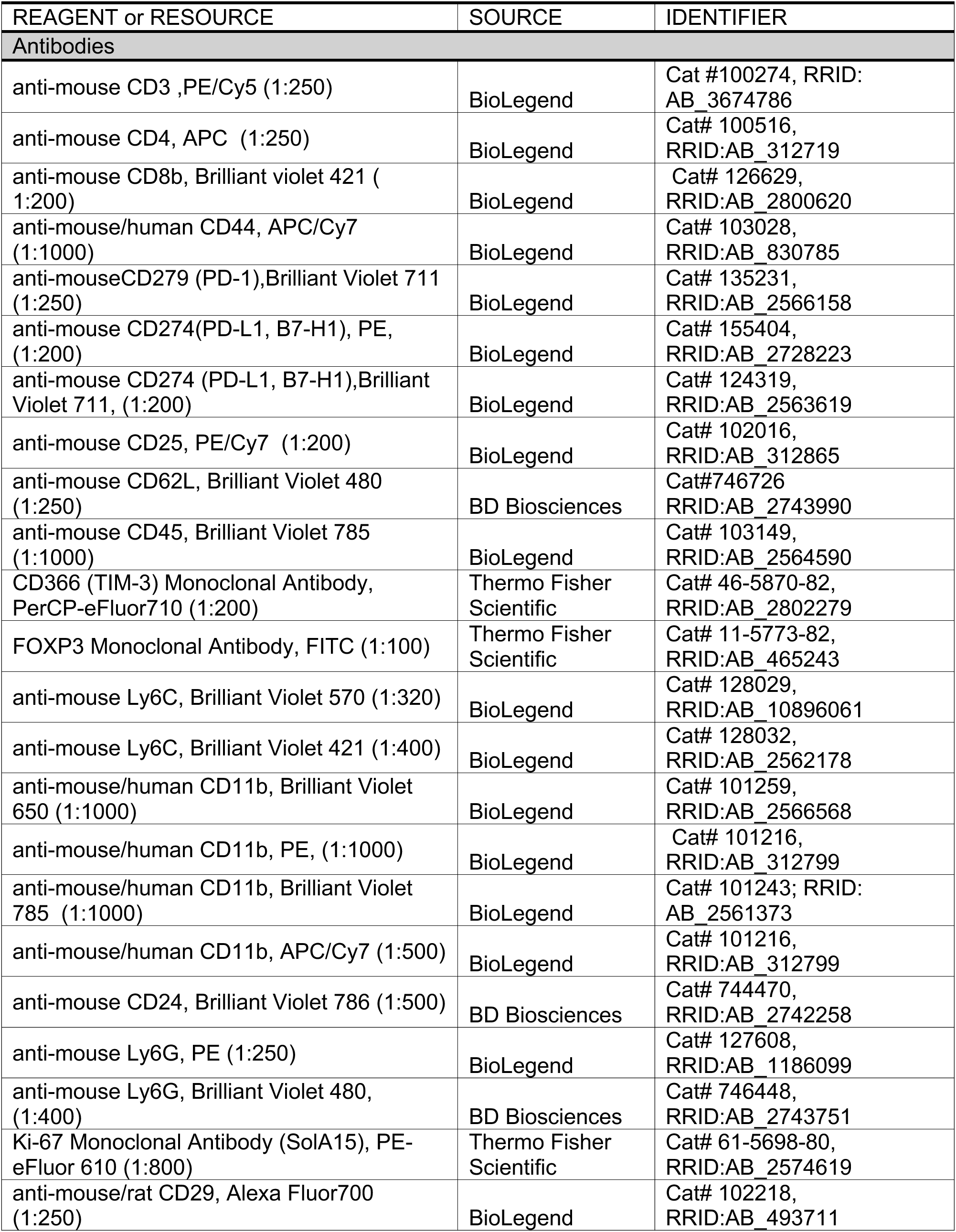
CyTOF panel for extracellular markers related to Figure 4J.

**Table 2:**
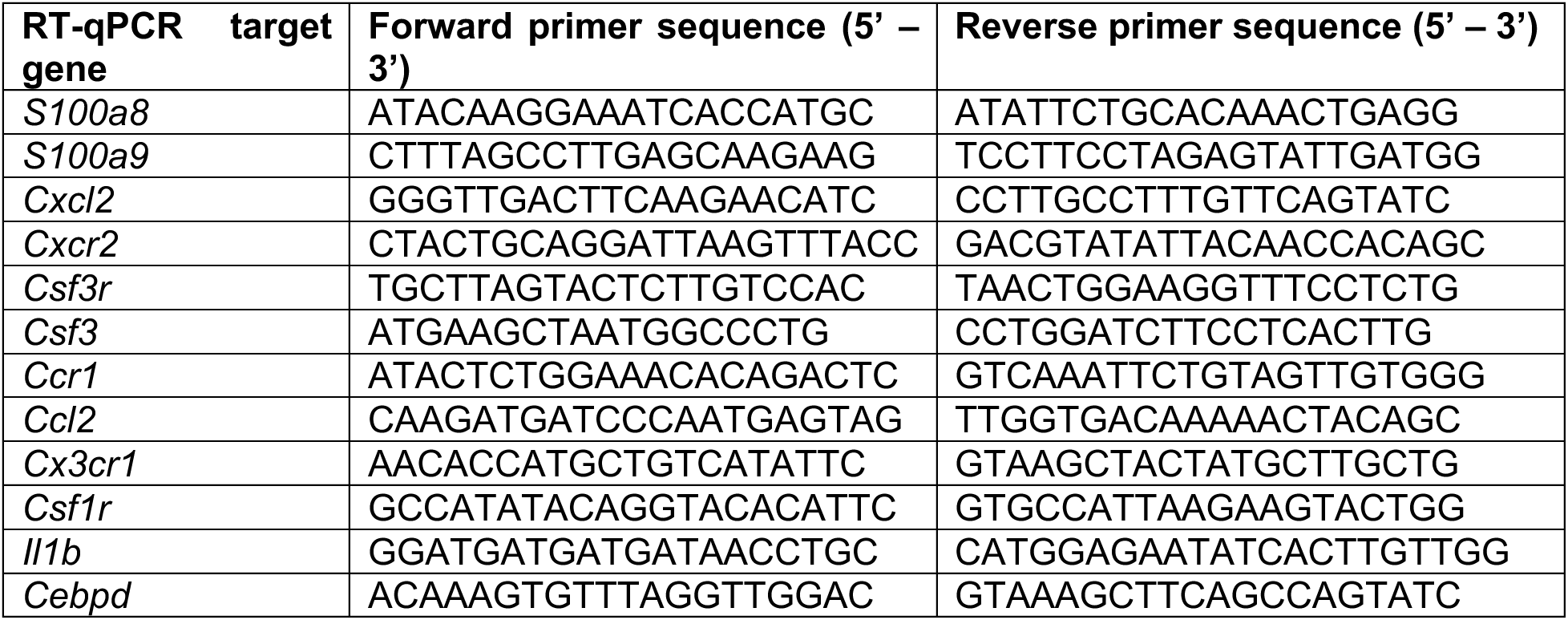
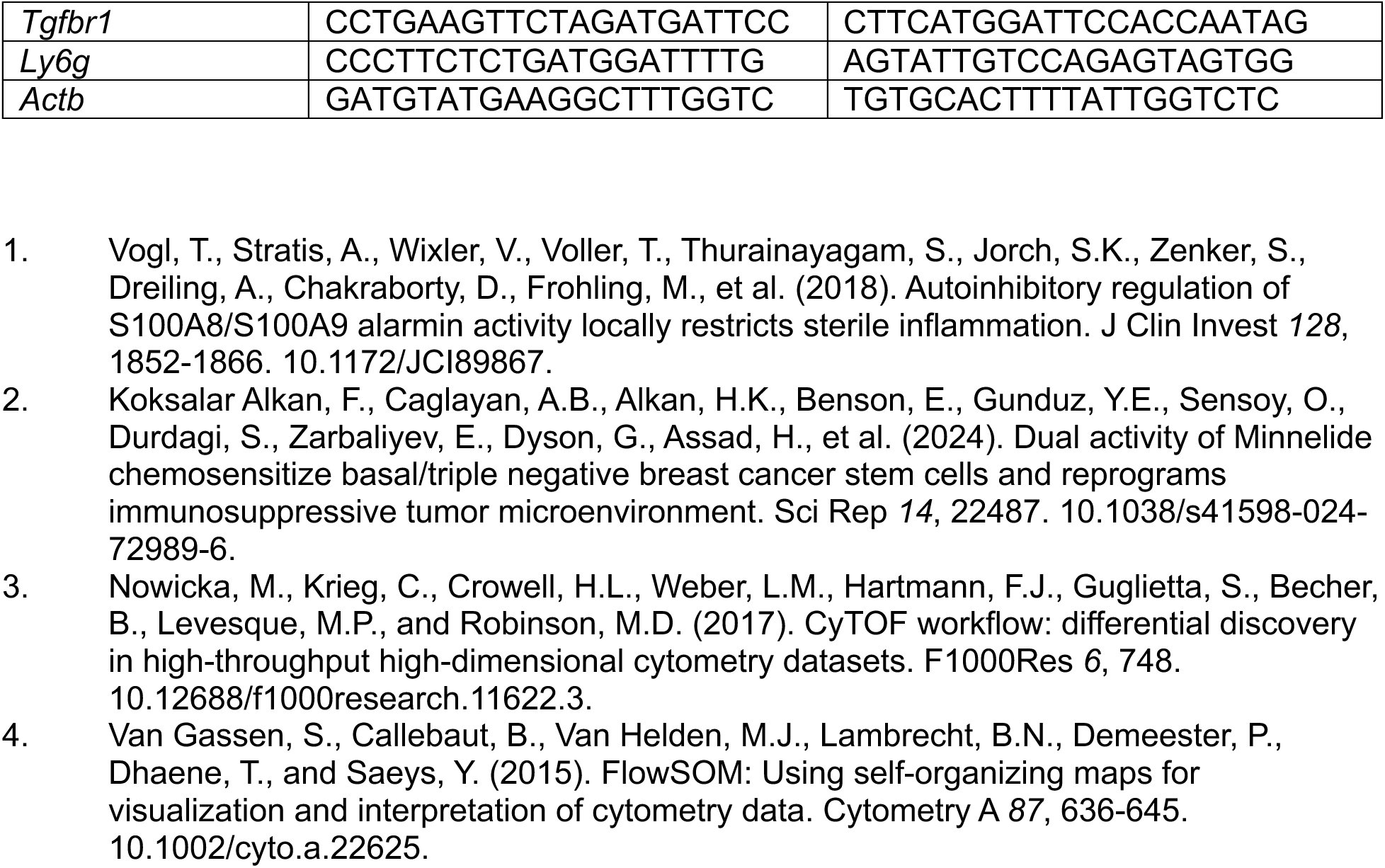
Oligonucleotides related to Figure1G, Figure 3P-R, Figure 4F and Figure 7H.

### Immunohistochemistry (IHC) and immunofluorescence (IF) staining

Paraffin embedded tissues were sectioned on positive charged slides with 5um thickness by the histology core facility. For IHC, slides were baked for 1 hour at 60°C. After washing with xylene, slides were passed through ethanol gradient, rinsed with distilled water and then with PBS. For antigen retrieval, slides were incubated with antigen retrieval solution (Citrate buffer pH 6) in the pressure cooker at the “high” setting for 10 minutes, then cooled down to the room temperature for staining. Slides were incubated in in 0.1% Triton X-100 in TBS for 15 min at room temperature, washed three times with PBS, and incubated with blocking solution TBS-T(TBS+0.1% Tween 20) supplemented with 10% serum of choice (BSA or Goat serum) for 1 hour at room temperature. Primary antibodies of CD15, Ki67, calprotectin were incubated in antibody solution TBST with 2% serum according to the instructions provided in the antibody datasheets overnight at 4°C or 4-6 hours at room temperature. Slides rinsed with TBS-T and incubated in secondary antibody solution with 2% serum according to the instructions provided in the antibody datasheets for 1 hour at room temperature. Slides were rinsed with PBS and mounting media used for glass coverslips. For IF, after antigen retrieval at 95°C for 10 minutes as performed in IHC, slides were incubated in 0.5%Triton X-100 in PBS for 15 min at room temperature and washed three times with PBS. Slides were incubated with blocking solution PBS-T (PBS+0.1% Tween 20) supplemented with 10% serum of choice (BSA or Goat serum) for 1 hour at room temperature. Primary antibodies were incubated in PBS-T with 2% antibodies overnight at 4°C or 4-6 hours at room temperature. Slides rinsed with PBS-T and incubated in secondary antibody solution with 2% serum according to the instructions provided in the antibody datasheets 1 hour at room temperature Slides were rinsed with PBS and mounting media with DAPI used for glass coverslips.

### Time-of-flight mass spectrometry (CyTOF)

Ex vivo spleen and tumors reduced to the single cell level as described above. Briefly, spleens were smashed with the white side of the syringe plunger through a 70μm sterile strainer into a 50 ml falcon tube and washed with magnesium and calcium free 1X DPBS (Gibco) without EDTA. Tumors were dissociated to the single cell level and washed through a 70μm sterile strainer into a 50ml falcon tub awith magnesium and calcium free 1X DPBS without EDTA. Red blood cells were eliminated by the ACK Lysing buffer (Gibco) according to manufacturers’ instructions. Single cells were submitted for the CyTOF labeling in 0.4% FBS containing D-PBS.

### Single-cell RNA sequencing (ScRNASeq)

Ex vivo spleen and tumors reduced to the single cell level as described above. Briefly, spleens were smashed with the white side of the syringe plunger through a 70μm sterile strainer into a 50 ml falcon tube and washed with magnesium and calcium free 1X DPBS (Gibco). Tumors were dissociated to the single cell level and washed through a 70μm sterile strainer into a 50ml falcon tub with magnesium and calcium free 1X DPBS without EDTA. Red blood cells were eliminated by the ACK Lysing buffer (Gibco) according to manufacturers’ instructions. Single cells were submitted to a genomics core in 0.2% BSA containing D-PBS. The sequencing libraries were constructed using 10X Genomics Chromium Single Cell Gene Expression 3’ Reagent kit v3 and sequenced using NextSeq500 sequencer from Illumina.

## QUATIFICATION AND STATISTICAL ANALYSIS

### Time-of-flight mass spectrometry (CyTOF) data processing

CyTOF data were analyzed using the CATALYST R package^44^. Signal intensities were normalized using an arcsinh transformation with a cofactor of 5. Live and singlet CD45+ cells were used for downstream analysis, and major immune populations were identified through unsupervised clustering with the FlowSOM^45^ algorithm using default parameters, followed by annotation based on canonical lineage marker expression. The optimal clustering resolution was determined using consensus clustering across k values ranging from 2 to 30, selecting the point where further increases produced minimal improvement in the cumulative distribution function (CDF) area. Median marker expression across clusters was visualized using heatmaps, and cellular relationships were projected using UMAP dimensionality reduction. Cell population frequencies were quantified for each sample and displayed using bar plots or box plots.

### Single-cell RNA sequencing data processing

The sequencing data were first processed using the 10X Genomics cellranger v6.1 aggr pipeline. The filtered count matrix was used as inputs and analyzed using the R package Seurat v5.0. Quality control measures were implemented in Seurat to remove cells with fewer less than 250 genes or more than 20% of mitochondria gene content. After normalization and scaling with SCTransform, samples were integrated using Seurat Canonical Correlation Analysis (CCA) integration, followed by dimensionality reduction and clustering. The top differentially expressed marker genes for each cluster were identified by the Seurat FindAllMarkers function. Cell types were predicted using the Cluster Identity Predictor (CIPR) package and annotated based on the known marker genes of major cell types. The spleen and tumor samples were analyzed separately. The neutrophils and myeloid progenitors from the tumor and spleen were re-integrated using Harmony and analyzed in the same way as described above. Cellular potency analysis of tumor epithelial cells was performed using the RunCytoTRACE function of sthe cop v0.8.9 package, which is a wrapper function that perform Predicts cellular developmental potential from single-cell RNA-seq data with the CytoTRACE 2 algorithm. Graphs were generated using plotting functions in the Seurat, scop and scCustomize v3.01 packages. Heatmaps were generated with ComplexHeatmap v2.22.0.

### Neutrophil trajectory analysis

Pseudotime trajectory analysis was performed with the assistance of Biomni (Phylo), an AI-powered scientific research platform. Biomni was used to convert Seurat objects to the AnnData format, construct neighborhood graphs on Harmony-corrected embeddings, run PAGA and diffusion pseudotime analyses (scanpy v1.11.4), generate publication-quality figures, and perform Mann-Whitney U and Fisher’s exact statistical tests. All analytical parameters, biological interpretations, and conclusions were reviewed and approved by the authors. The analysis and figure generation scripts were reviewed by experts and tested on a local machine, and reproducible results were confirmed.

*Statement about AI tool use:* The authors used Biomni (Phylo), an AI-assisted computational biology platform, to support data processing, cell type annotation, trajectory analysis, and figure generation. The authors take full responsibility for the integrity of the data and the accuracy of the analysis.

### Statistical analysis

GraphPad (Prism) software was used for the statistical analysis of *in vivo* and *ex vivo* bioluminescent imaging, *ex vivo* tissues, *in vitro* and *in vivo* flow cytometry and RT-qPCR data. Data was represented as mean ± standard error of the means (SEM). Statistical methods applied between two groups were two-tailed Student *t*-test and 3 or more groups were one-way ANOVA followed by Tukey’s multiple comparison test. P-values are reported as n: * = p % 0.05, ** = p % 0.01, *** = p % 0.001, **** = p % 0.0001. Detailed statistical methods and sample size (n) of animals/tissues/cells were indicated in the figure legends.

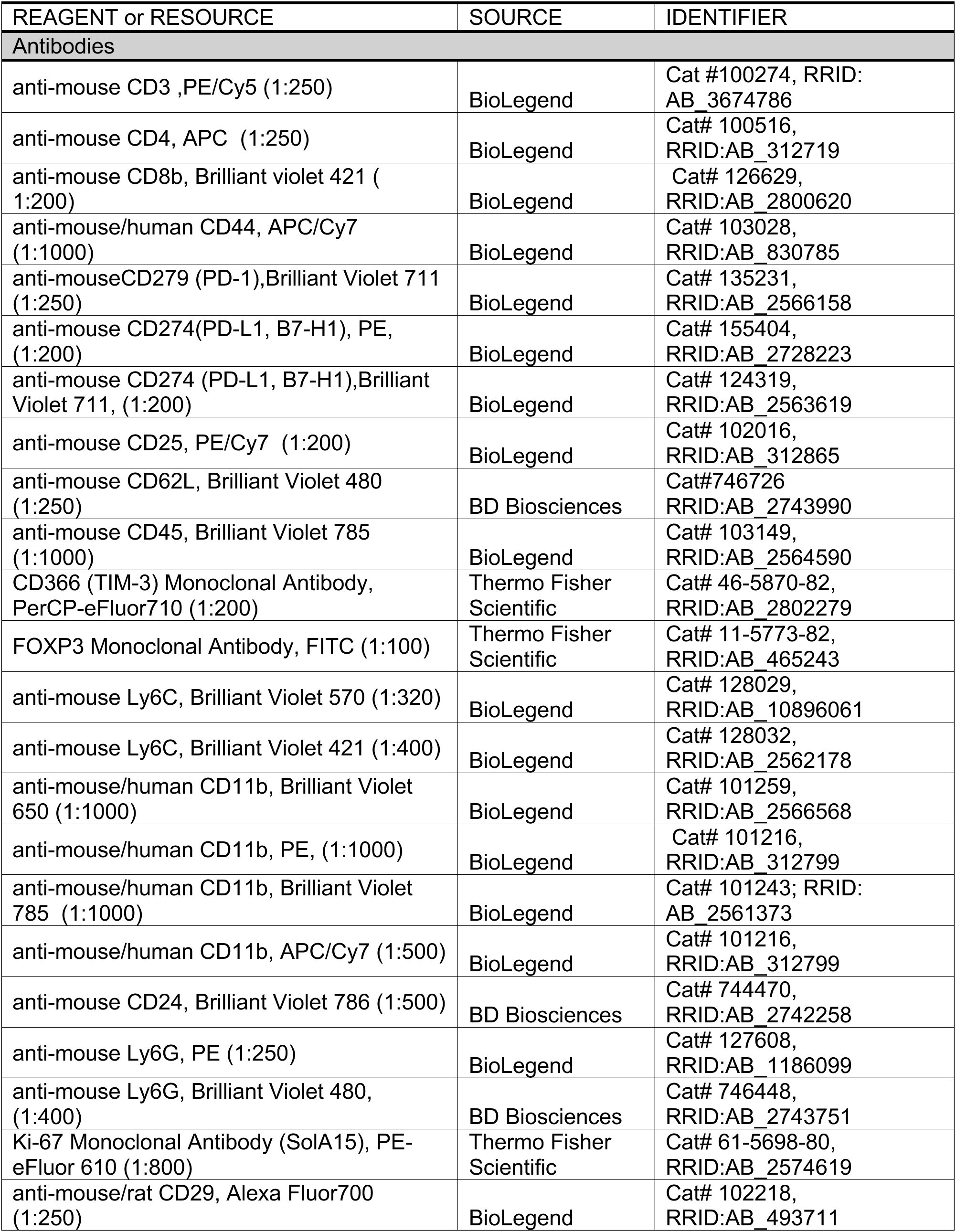
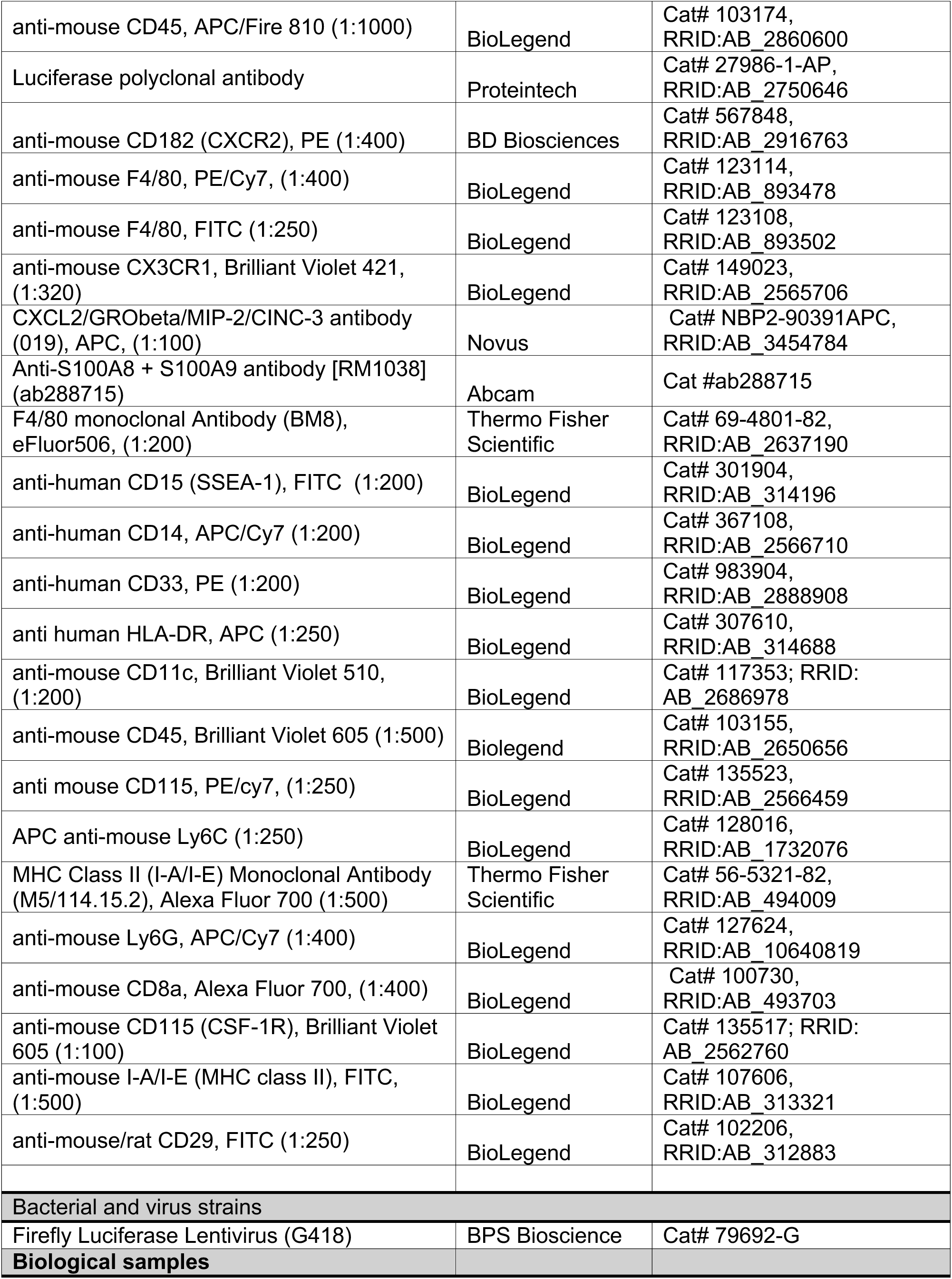
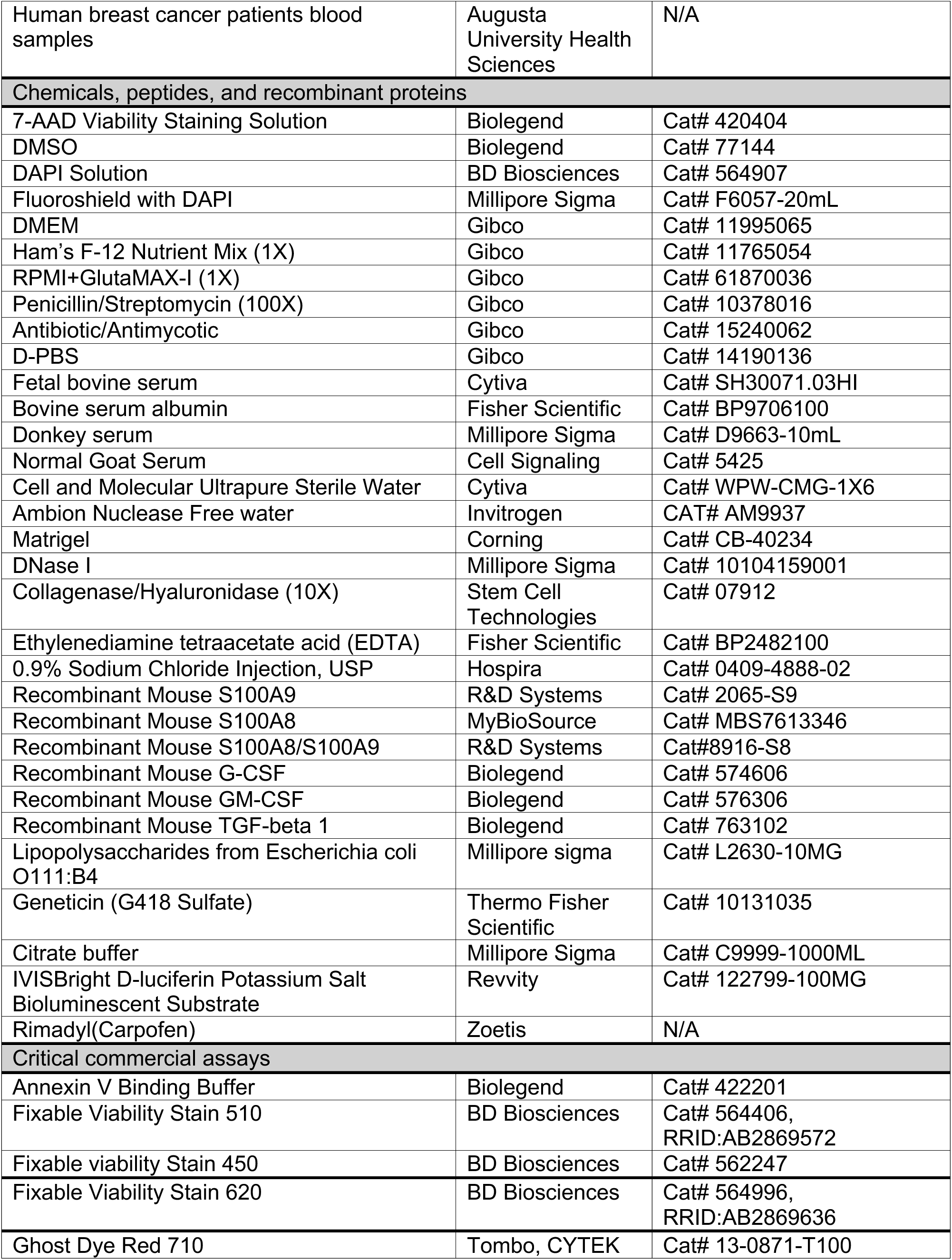
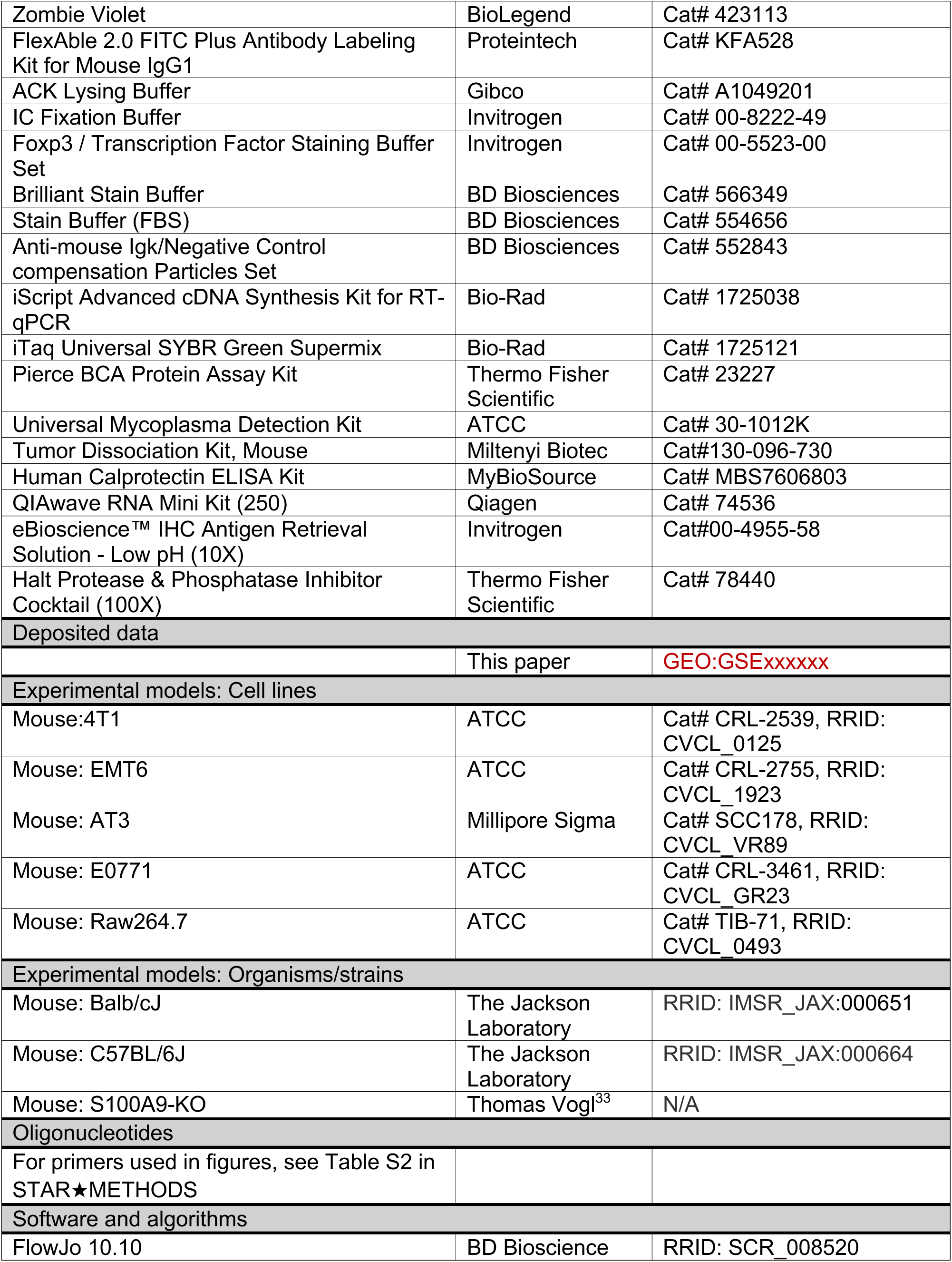
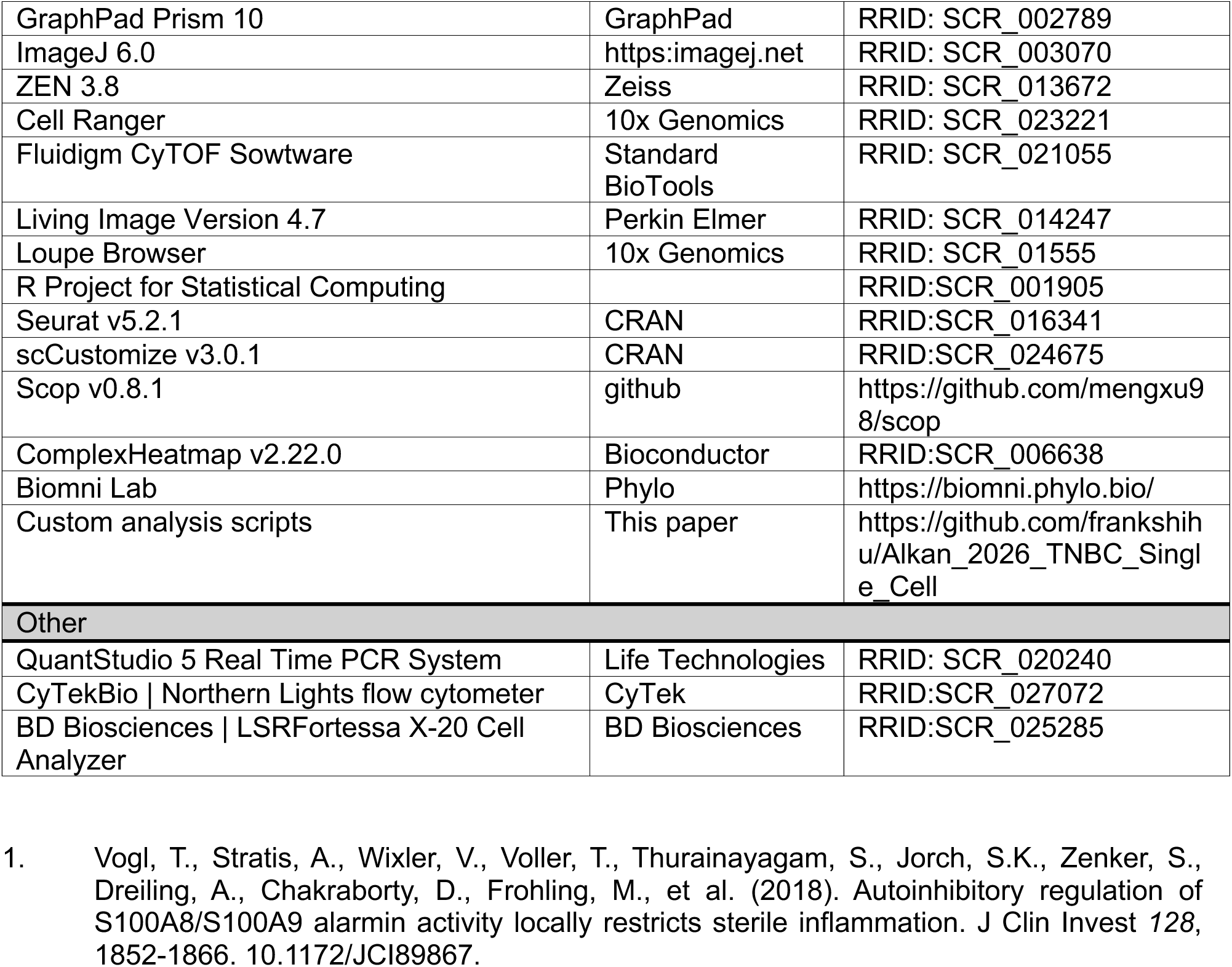
Key resources table.

## Supplemental Figure Legends

**Figure S1.** Metastatic 4T1 tumors exhibit accelerated growth and systemic expansion compared to non-invasive EMT6 tumors. (**A-B**) Representative images 5 independent tumors and spleens of EMT6 and 4T1 tumor bearing mice used and pooled for scRNA-Seq analyses. One-Way ANOVA mean ± SEM test applied to compare spleen sizes *p **<**0.05, **p **<**0.01. **(C)** Representative images of excised primary tumors from EMT6 and 4T1 tumor-bearing mice at weeks 1-3, demonstrating accelerated tumor growth in 4T1 relative to EMT6. These tissues were used for immune profiling using high parameter spectral flow cytometry. **(D)** Representative images and quantification of spleens from non–tumor-bearing (Non-T), EMT6, and 4T1 mice, showing progressive splenomegaly in 4T1 tumor bearing animals. **(E)** Quantification of ex vivo tumor weights of 4T1 and EMT6 tumor-bearing mice over time. **F)** Quantification of spleen weight during the three week time course of tumor growth, indicating systemic immune expansion and splenomegaly in 4T1 tumor bearing mice compared to EMT6 tumor bearing mice. Data represent mean ± SEM; statistical significance determined Welch-t-test (*p **<**0.05, **p **<**0.01, *** p **<** 0.005, **** p **<** 0.001). **(G)** Bioluminescence imaging (BLI) of lungs during the three week time course of tumor growth, showing relatively higher metastatic burden in 4T1 compared to EMT6 tumor bearing mice.

**Figure S2.** Transcriptional clustering reveals distinct myeloid and lymphoid immune programs in TNBC tumors. Heatmap showing hierarchical clustering of differentially expressed genes across tumor-infiltrating immune cell populations, including B cells, CD4⁺ and CD8⁺ T cells, regulatory T cells, NK cells, monocytes, macrophages, dendritic cells, and neutrophil subsets. Granulocytic populations cluster together and exhibit enrichment of inflammatory and S100A8/A9-associated gene programs (Ly6G+_Neutrophil, Il1b.hu_Neutrophil, and Ccl3.hi_Neutrophil), whereas lymphoid populations segregate into distinct clusters characterized by activation and adaptive immune signatures. Expression values are z-score normalized.

**Figure S3.** Monocyte/macrophage targeting in EMT6 tumors minimally impacts systemic immunity and anti-tumor memory. **(A)** Flow cytometric analysis of monocyte/macrophage populations across tumor, PBMC, spleen, and lung compartments in EMT6 tumor-bearing mice. **(B)** In vivo imaging and tumor growth analysis following anti-CD115–mediated depletion of monocyte/macrophage populations. **(C)** Immune profiling of PBMCs and tumors showing modest changes in myeloid populations upon CD115 targeting. **(D)** Tumor resection and re-challenge experiments demonstrating preserved anti-tumor immunity and memory responses in EMT6 models despite monocyte/macrophage depletion. **(E)** Quantification of immune subsets and tumor burden across treatment conditions. These findings indicate that, in contrast to neutrophil-driven suppression in metastatic tumors, monocyte/macrophage populations in EMT6 tumors do not drive systemic immune dysfunction.

**Figure S4.** PD-L1 blockade in WT animals reprograms systemic immune composition without showing any effect on tumor growth and metastatic formation in the lungs. **(A)** Surface PD-L1 expression across tumor models as measured by flow cytometry (n= each) (Data represent mean ± SEM; statistical significance determined Welch-t-test (*p **<**0.05) **(B-C)** Tumor growth kinetics measured by in vivo BLI in WT C57BL/6 mice treated with Isotype or Anti-PDL1 (n=5 each). **(D)** Spleen weights compared between WT C57BL/6 mice treated with Isotype or Anti-PDL1 (Data represent mean ± SEM; statistical significance determined Welch-t-test (n.s; non-significant). **(E)** Ex vivo lung BLI quantification showing reduced metastatic burden in anti-PDL1 treated mice. **(F-G)** Flow cytometric analysis of tumor and splenic immune populations demonstrating reduced CXCR2+ CD11b+ and Ly6G+Ly6C– neutrophils and increased CX3CR1 representing monocyte/macrophage populations in anti-PDL1 treated mice compared to isotype controls (n=4 each). (Data represent mean ± SEM; statistical significance determined Welch-t-test (*p **<**0.05).

---

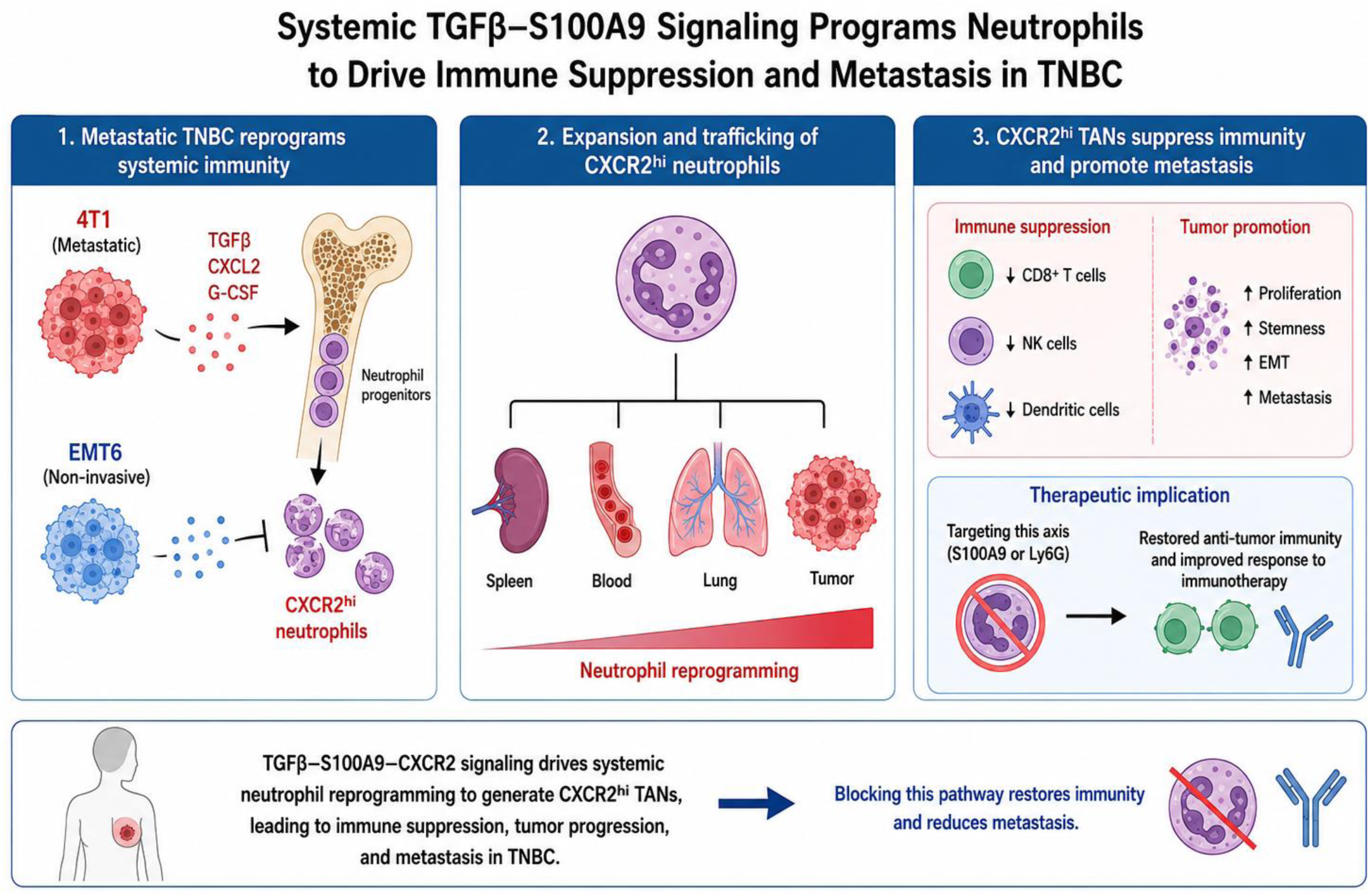

## REFERENCES

1. Rakha, E.A., Elsheikh, S.E., Aleskandarany, M.A., Habashi, H.O., Green, A.R., Powe, D.G., El-Sayed, M.E., Benhasouna, A., Brunet, J.S., Akslen, L.A., et al. (2009). Triple-negative breast cancer: distinguishing between basal and nonbasal subtypes. Clin Cancer Res 15, 2302–2310. 1078-0432.CCR-08-2132 [pii] 10.1158/1078-0432.CCR-08-2132.

2. Dent, R., Trudeau, M., Pritchard, K.I., Hanna, W.M., Kahn, H.K., Sawka, C.A., Lickley, L.A., Rawlinson, E., Sun, P., and Narod, S.A. (2007). Triple-negative breast cancer: clinical features and patterns of recurrence. Clin Cancer Res 13, 4429–4434. 13/15/4429 [pii] 10.1158/1078-0432.CCR-06-3045.

3. Keren, L., Bosse, M., Marquez, D., Angoshtari, R., Jain, S., Varma, S., Yang, S.R., Kurian, A., Van Valen, D., West, R., et al. (2018). A Structured Tumor-Immune Microenvironment in Triple Negative Breast Cancer Revealed by Multiplexed Ion Beam Imaging. Cell 174, 1373–1387 e1319. 10.1016/j.cell.2018.08.039.

4. Bareche, Y., Buisseret, L., Gruosso, T., Girard, E., Venet, D., Dupont, F., Desmedt, C., Larsimont, D., Park, M., Rothe, F., et al. (2020). Unraveling Triple-Negative Breast Cancer Tumor Microenvironment Heterogeneity: Towards an Optimized Treatment Approach. J Natl Cancer Inst 112, 708–719. 10.1093/jnci/djz208.

5. Ugel, S., De Sanctis, F., Mandruzzato, S., and Bronte, V. (2015). Tumor-induced myeloid deviation: when myeloid-derived suppressor cells meet tumor-associated macrophages. J Clin Invest 125, 3365–3376. 10.1172/JCI80006.

6. Awad, R.M., De Vlaeminck, Y., Maebe, J., Goyvaerts, C., and Breckpot, K. (2018). Turn Back the TIMe: Targeting Tumor Infiltrating Myeloid Cells to Revert Cancer Progression. Frontiers in immunology 9, 1977. 10.3389/fimmu.2018.01977.

7. Bronte, V., Brandau, S., Chen, S.H., Colombo, M.P., Frey, A.B., Greten, T.F., Mandruzzato, S., Murray, P.J., Ochoa, A., Ostrand-Rosenberg, S., et al. (2016). Recommendations for myeloid-derived suppressor cell nomenclature and characterization standards. Nat Commun 7, 12150. 10.1038/ncomms12150.

8. Hiam-Galvez, K.J., Allen, B.M., and Spitzer, M.H. (2021). Systemic immunity in cancer. Nat Rev Cancer 21, 345–359. 10.1038/s41568-021-00347-z.

9. Casbon, A.J., Reynaud, D., Park, C., Khuc, E., Gan, D.D., Schepers, K., Passegue, E., and Werb, Z. (2015). Invasive breast cancer reprograms early myeloid differentiation in the bone marrow to generate immunosuppressive neutrophils. Proc Natl Acad Sci U S A 112, E566–575. 10.1073/pnas.1424927112.

10. Templeton, A.J., McNamara, M.G., Seruga, B., Vera-Badillo, F.E., Aneja, P., Ocana, A., Leibowitz-Amit, R., Sonpavde, G., Knox, J.J., Tran, B., et al. (2014). Prognostic role of neutrophil-to-lymphocyte ratio in solid tumors: a systematic review and meta-analysis. J Natl Cancer Inst 106, dju124. 10.1093/jnci/dju124.

11. Piranlioglu, R., Lee, E., Ouzounova, M., Bollag, R.J., Vinyard, A.H., Arbab, A.S., Marasco, D., Guzel, M., Cowell, J.K., Thangaraju, M., et al. (2019). Primary tumor-induced immunity eradicates disseminated tumor cells in syngeneic mouse model. Nat Commun 10, 1430. 10.1038/s41467-019-09015-1.

12. Ouzounova, M., Lee, E., Piranlioglu, R., El Andaloussi, A., Kolhe, R., Demirci, M.F., Marasco, D., Asm, I., Chadli, A., Hassan, K.A., et al. (2017). Monocytic and granulocytic myeloid derived suppressor cells differentially regulate spatiotemporal tumour plasticity during metastatic cascade. Nat Commun 8, 14979. 10.1038/ncomms14979.

13. Kaur, P., Nagaraja, G.M., Zheng, H., Gizachew, D., Galukande, M., Krishnan, S., and Asea, A. (2012). A mouse model for triple-negative breast cancer tumor-initiating cells (TNBC-TICs) exhibits similar aggressive phenotype to the human disease. BMC Cancer 12, 120. 10.1186/1471-2407-12-120.

14. Bajrami, B., Zhu, H., Kwak, H.J., Mondal, S., Hou, Q., Geng, G., Karatepe, K., Zhang, Y.C., Nombela-Arrieta, C., Park, S.Y., et al. (2016). G-CSF maintains controlled neutrophil mobilization during acute inflammation by negatively regulating CXCR2 signaling. J Exp Med 213, 1999–2018. 10.1084/jem.20160393.

15. Huang, M., Wu, R., Chen, L., Peng, Q., Li, S., Zhang, Y., Zhou, L., and Duan, L. (2019). S100A9 Regulates MDSCs-Mediated Immune Suppression via the RAGE and TLR4 Signaling Pathways in Colorectal Carcinoma. Frontiers in immunology 10, 2243. 10.3389/fimmu.2019.02243.

16. Engblom, C., Pfirschke, C., and Pittet, M.J. (2016). The role of myeloid cells in cancer therapies. Nat Rev Cancer 16, 447–462. 10.1038/nrc.2016.54.

17. Binnewies, M., Roberts, E.W., Kersten, K., Chan, V., Fearon, D.F., Merad, M., Coussens, L.M., Gabrilovich, D.I., Ostrand-Rosenberg, S., Hedrick, C.C., et al. (2018). Understanding the tumor immune microenvironment (TIME) for effective therapy. Nat Med 24, 541–550. 10.1038/s41591-018-0014-x.

18. Jauch-Speer, S.L., Herrera-Rivero, M., Ludwig, N., Veras De Carvalho, B.C., Martens, L., Wolf, J., Imam Chasan, A., Witten, A., Markus, B., Schieffer, B., et al. (2022). C/EBPdelta-induced epigenetic changes control the dynamic gene transcription of S100a8 and S100a9. Elife 11. 10.7554/eLife.75594.

19. Vogl, T., Leukert, N., Barczyk, K., Strupat, K., and Roth, J. (2006). Biophysical characterization of S100A8 and S100A9 in the absence and presence of bivalent cations. Biochim Biophys Acta 1763, 1298–1306. 10.1016/j.bbamcr.2006.08.028.

20. Mousset, A., Lecorgne, E., Bourget, I., Lopez, P., Jenovai, K., Cherfils-Vicini, J., Dominici, C., Rios, G., Girard-Riboulleau, C., Liu, B., et al. (2023). Neutrophil extracellular traps formed during chemotherapy confer treatment resistance via TGF-beta activation. Cancer Cell 41, 757–775 e710. 10.1016/j.ccell.2023.03.008.

21. Pang, Y., Gara, S.K., Achyut, B.R., Li, Z., Yan, H.H., Day, C.P., Weiss, J.M., Trinchieri, G., Morris, J.C., and Yang, L. (2013). TGF-beta signaling in myeloid cells is required for tumor metastasis. Cancer Discov 3, 936–951. 10.1158/2159-8290.CD-12-0527.

22. Klughammer, J., Abravanel, D.L., Segerstolpe, A., Blosser, T.R., Goltsev, Y., Cui, Y., Goodwin, D.R., Sinha, A., Ashenberg, O., Slyper, M., et al. (2024). A multi-modal single-cell and spatial expression map of metastatic breast cancer biopsies across clinicopathological features. Nat Med 30, 3236–3249. 10.1038/s41591-024-03215-z.

23. Alshetaiwi, H., Pervolarakis, N., McIntyre, L.L., Ma, D., Nguyen, Q., Rath, J.A., Nee, K., Hernandez, G., Evans, K., Torosian, L., et al. (2020). Defining the emergence of myeloid-derived suppressor cells in breast cancer using single-cell transcriptomics. Sci Immunol 5. 10.1126/sciimmunol.aay6017.

24. Winkler, J., Tan, W., Diadhiou, C.M., McGinnis, C.S., Abbasi, A., Hasnain, S., Durney, S., Atamaniuc, E., Superville, D., Awni, L., et al. (2024). Single-cell analysis of breast cancer metastasis reveals epithelial-mesenchymal plasticity signatures associated with poor outcomes. J Clin Invest 134. 10.1172/JCI164227.

25. Hanna, R.N., Cekic, C., Sag, D., Tacke, R., Thomas, G.D., Nowyhed, H., Herrley, E., Rasquinha, N., McArdle, S., Wu, R., et al. (2015). Patrolling monocytes control tumor metastasis to the lung. Science 350, 985–990. 10.1126/science.aac9407.

26. Yue, Y., Zhang, Q., and Sun, Z. (2021). CX3CR1 Acts as a Protective Biomarker in the Tumor Microenvironment of Colorectal Cancer. Frontiers in immunology 12, 758040. 10.3389/fimmu.2021.758040.

27. Yamauchi, T., Hoki, T., Oba, T., Jain, V., Chen, H., Attwood, K., Battaglia, S., George, S., Chatta, G., Puzanov, I., et al. (2021). T-cell CX3CR1 expression as a dynamic blood-based biomarker of response to immune checkpoint inhibitors. Nat Commun 12, 1402. 10.1038/s41467-021-21619-0.

28. Abdelfatah, E., Long, M.D., Kajihara, R., Oba, T., Yamauchi, T., Chen, H., Sarkar, J., Attwood, K., Matsuzaki, J., Segal, B.H., et al. (2023). Predictive and Prognostic Implications of Circulating CX3CR1(+) CD8(+) T Cells in Non-Small Cell Lung Cancer Patients Treated with Chemo-Immunotherapy. Cancer Res Commun 3, 510–520. 10.1158/2767-9764.CRC-22-0383.

29. Stephan, J.K., Knerr, T., Wells, C.K., Gu, Z., Johnson, S., Jobe, T.K., Isaacs, W.S., Hill, B.G., and Wysoczynski, M. (2025). G-CSF-Induced Emergency Granulopoiesis Modulates Neutrophil Effector Function in Mice. Stem Cell Rev Rep 21, 1113–1126. 10.1007/s12015-025-10885-w.

30. Kwak, T., Wang, F., Deng, H., Condamine, T., Kumar, V., Perego, M., Kossenkov, A., Montaner, L.J., Xu, X., Xu, W., et al. (2020). Distinct Populations of Immune-Suppressive Macrophages Differentiate from Monocytic Myeloid-Derived Suppressor Cells in Cancer. Cell reports 33, 108571. 10.1016/j.celrep.2020.108571.

31. Cheng, P., Corzo, C.A., Luetteke, N., Yu, B., Nagaraj, S., Bui, M.M., Ortiz, M., Nacken, W., Sorg, C., Vogl, T., et al. (2008). Inhibition of dendritic cell differentiation and accumulation of myeloid-derived suppressor cells in cancer is regulated by S100A9 protein. J Exp Med 205, 2235–2249. 10.1084/jem.20080132.

32. Garcia, V., Perera, Y.R., and Chazin, W.J. (2022). A Structural Perspective on Calprotectin as a Ligand of Receptors Mediating Inflammation and Potential Drug Target. Biomolecules 12. 10.3390/biom12040519.

33. Vogl, T., Stratis, A., Wixler, V., Voller, T., Thurainayagam, S., Jorch, S.K., Zenker, S., Dreiling, A., Chakraborty, D., Frohling, M., et al. (2018). Autoinhibitory regulation of S100A8/S100A9 alarmin activity locally restricts sterile inflammation. J Clin Invest 128, 1852–1866. 10.1172/JCI89867.

34. Schmid, P., Adams, S., Rugo, H.S., Schneeweiss, A., Barrios, C.H., Iwata, H., Dieras, V., Hegg, R., Im, S.A., Shaw Wright, G., et al. (2018). Atezolizumab and Nab-Paclitaxel in Advanced Triple-Negative Breast Cancer. N Engl J Med. 10.1056/NEJMoa1809615.

35. Bianchini, G., De Angelis, C., Licata, L., and Gianni, L. (2022). Treatment landscape of triple-negative breast cancer – expanded options, evolving needs. Nat Rev Clin Oncol 19, 91–113. 10.1038/s41571-021-00565-2.

36. Bakker, N.A.M., Garner, H., van Dyk, E., Champanhet, E., Klaver, C., Duijst, M., Voorwerk, L., Nederlof, I., Voorthuis, R., Liefaard, M.C., et al. (2025). Triple-negative breast cancer modifies the systemic immune landscape and alters neutrophil functionality. NPJ Breast Cancer 11, 5. 10.1038/s41523-025-00721-2.

37. Wu, Y., Ma, J., Yang, X., Nan, F., Zhang, T., Ji, S., Rao, D., Feng, H., Gao, K., Gu, X., et al. (2024). Neutrophil profiling illuminates anti-tumor antigen-presenting potency. Cell 187, 1422–1439 e1424. 10.1016/j.cell.2024.02.005.

38. Ng, M.S.F., Kwok, I., Tan, L., Shi, C., Cerezo-Wallis, D., Tan, Y., Leong, K., Calvo, G.F., Yang, K., Zhang, Y., et al. (2024). Deterministic reprogramming of neutrophils within tumors. Science 383, eadf6493. 10.1126/science.adf6493.

39. Salcher, S., Sturm, G., Horvath, L., Untergasser, G., Kuempers, C., Fotakis, G., Panizzolo, E., Martowicz, A., Trebo, M., Pall, G., et al. (2022). High-resolution single-cell atlas reveals diversity and plasticity of tissue-resident neutrophils in non-small cell lung cancer. Cancer Cell 40, 1503–1520 e1508. 10.1016/j.ccell.2022.10.008.

40. Garner, H., Martinovic, M., Liu, N.Q., Bakker, N.A.M., Velilla, I.Q., Hau, C.S., Vrijland, K., Kaldenbach, D., Kok, M., de Wit, E., and de Visser, K.E. (2025). Understanding and reversing mammary tumor-driven reprogramming of myelopoiesis to reduce metastatic spread. Cancer Cell 43, 1279–1295 e1279. 10.1016/j.ccell.2025.04.007.

41. Begg, L.R., Orriols, A.M., Zannikou, M., Yeh, C., Vadlamani, P., Kanojia, D., Bolin, R., Dunne, S.F., Balakrishnan, S., Camarda, R., et al. (2024). S100A8/A9 predicts response to PIM kinase and PD-1/PD-L1 inhibition in triple-negative breast cancer mouse models. Commun Med (Lond) 4, 22. 10.1038/s43856-024-00444-8.

42. Camargo, S., Moskowitz, O., Giladi, A., Levinson, M., Balaban, R., Gola, S., Raizman, A., Lipczyc, K., Richter, A., Keren-Khadmy, N., et al. (2025). Neutrophils physically interact with tumor cells to form a signaling niche promoting breast cancer aggressiveness. Nat Cancer 6, 540–558. 10.1038/s43018-025-00924-3.

43. Koksalar Alkan, F., Caglayan, A.B., Alkan, H.K., Benson, E., Gunduz, Y.E., Sensoy, O., Durdagi, S., Zarbaliyev, E., Dyson, G., Assad, H., et al. (2024). Dual activity of Minnelide chemosensitize basal/triple negative breast cancer stem cells and reprograms immunosuppressive tumor microenvironment. Scientific reports 14, 22487. 10.1038/s41598-024-72989-6.

44. Nowicka, M., Krieg, C., Crowell, H.L., Weber, L.M., Hartmann, F.J., Guglietta, S., Becher, B., Levesque, M.P., and Robinson, M.D. (2017). CyTOF workflow: differential discovery in high-throughput high-dimensional cytometry datasets. F1000Res 6, 748. 10.12688/f1000research.11622.3.

45. Van Gassen, S., Callebaut, B., Van Helden, M.J., Lambrecht, B.N., Demeester, P., Dhaene, T., and Saeys, Y. (2015). FlowSOM: Using self-organizing maps for visualization and interpretation of cytometry data. Cytometry A 87, 636–645. 10.1002/cyto.a.22625.

