## Supplemental Data for "S100A9-Dependent CXCR2^hi^ Neutrophils Mediate Systemic Immune Suppression and Checkpoint Resistance in Metastatic TNBC"

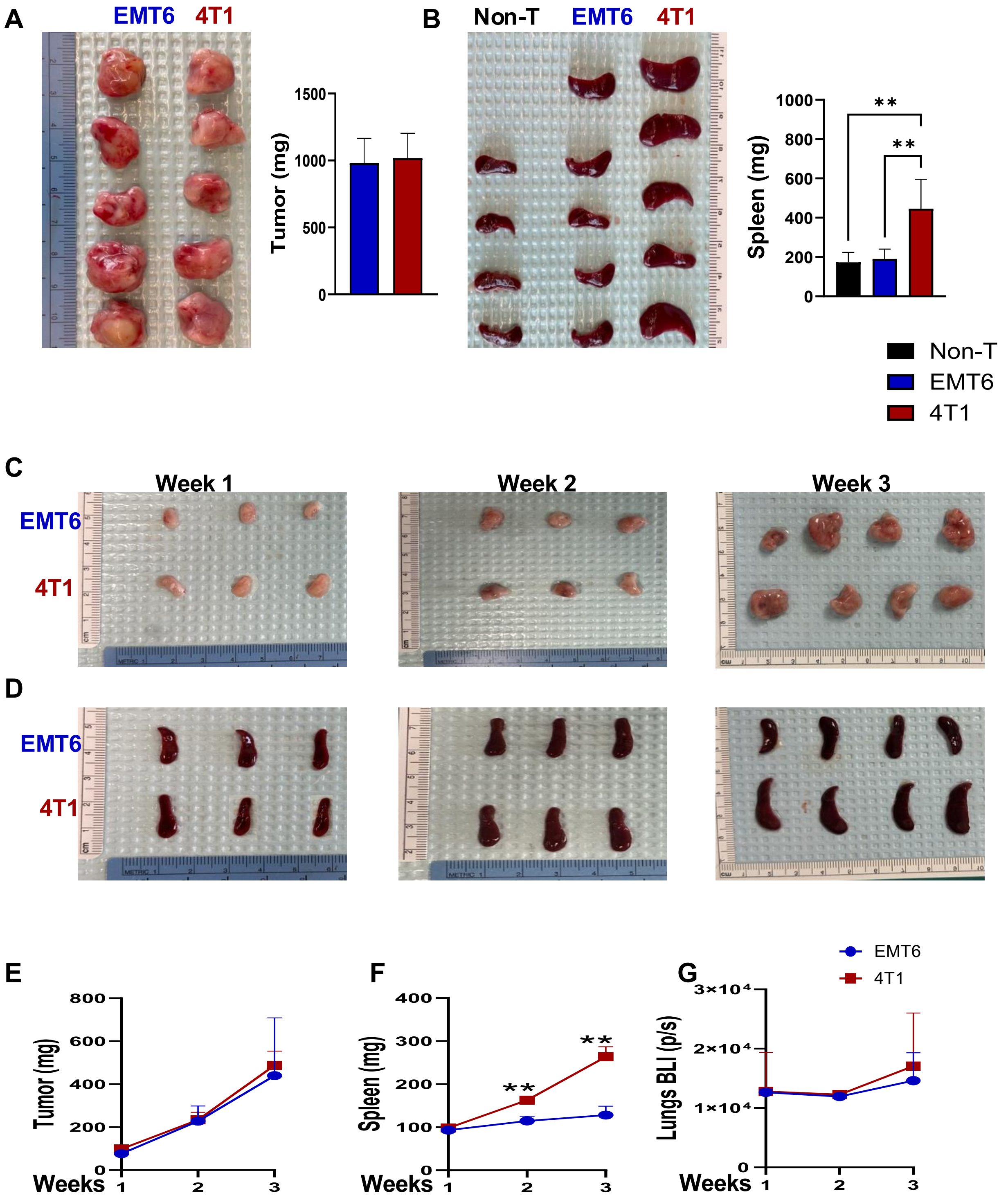


Supplemental Figure S1

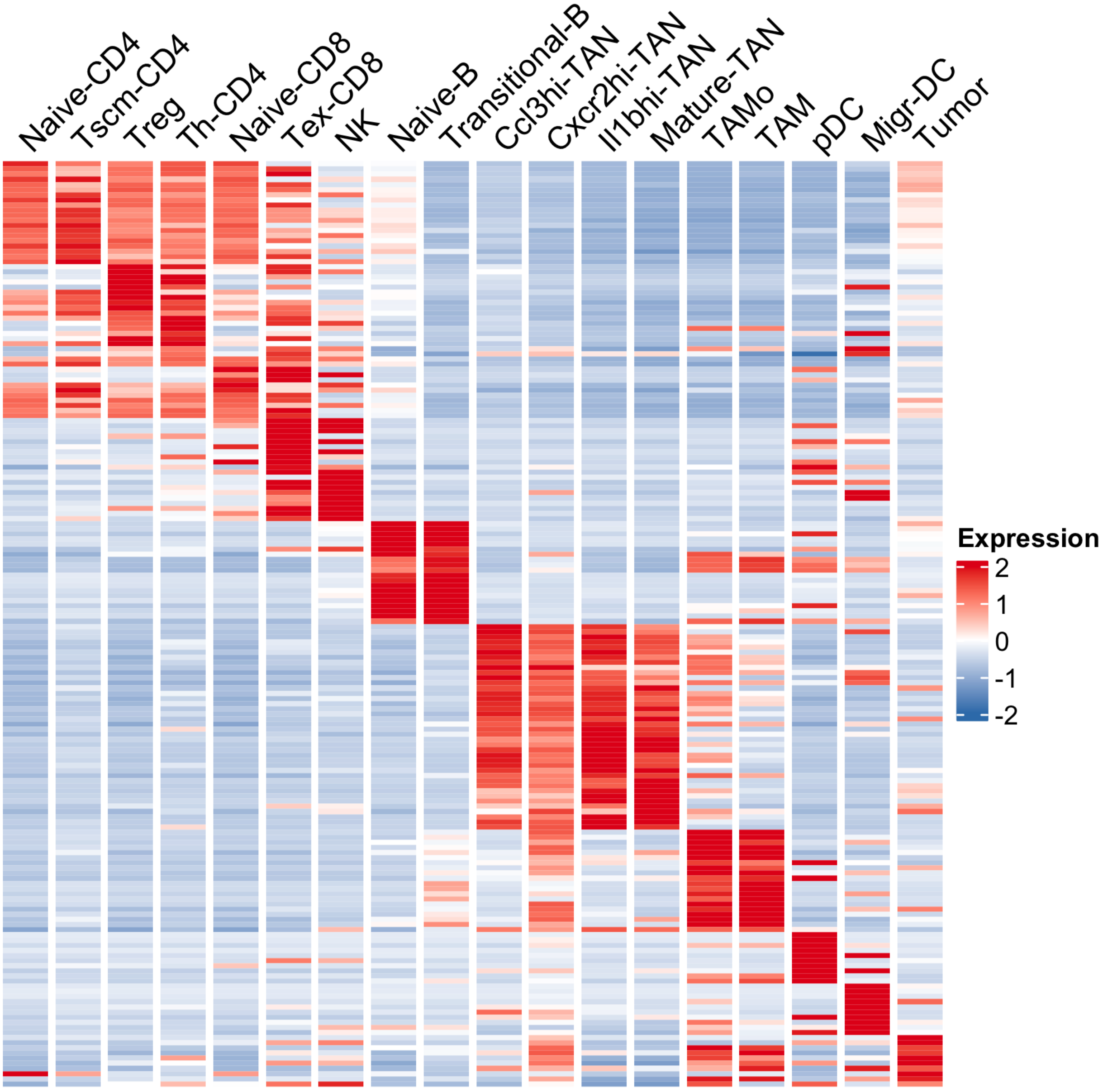


**Supplemental Figure S1**

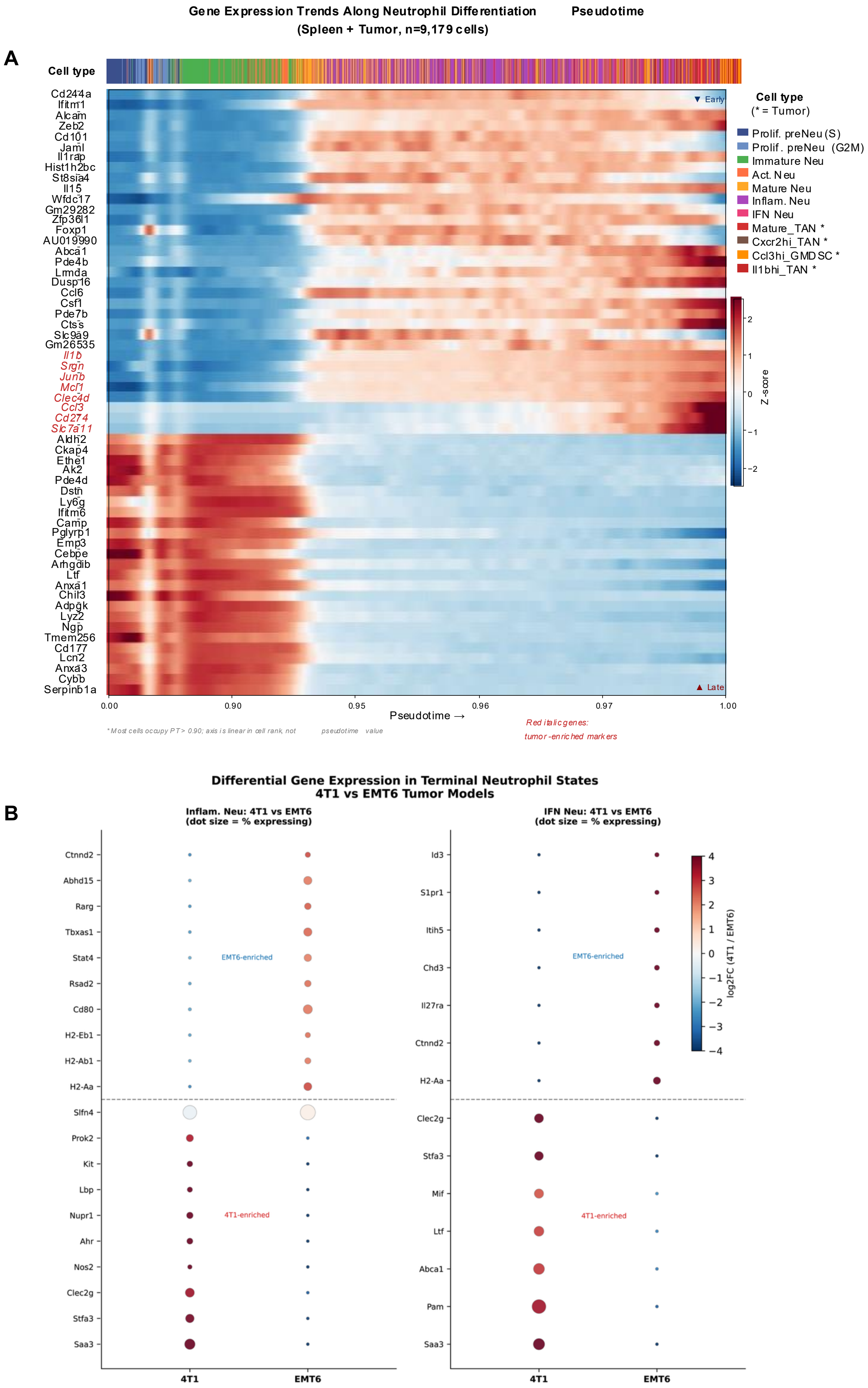


**Supplemental Figure S3**

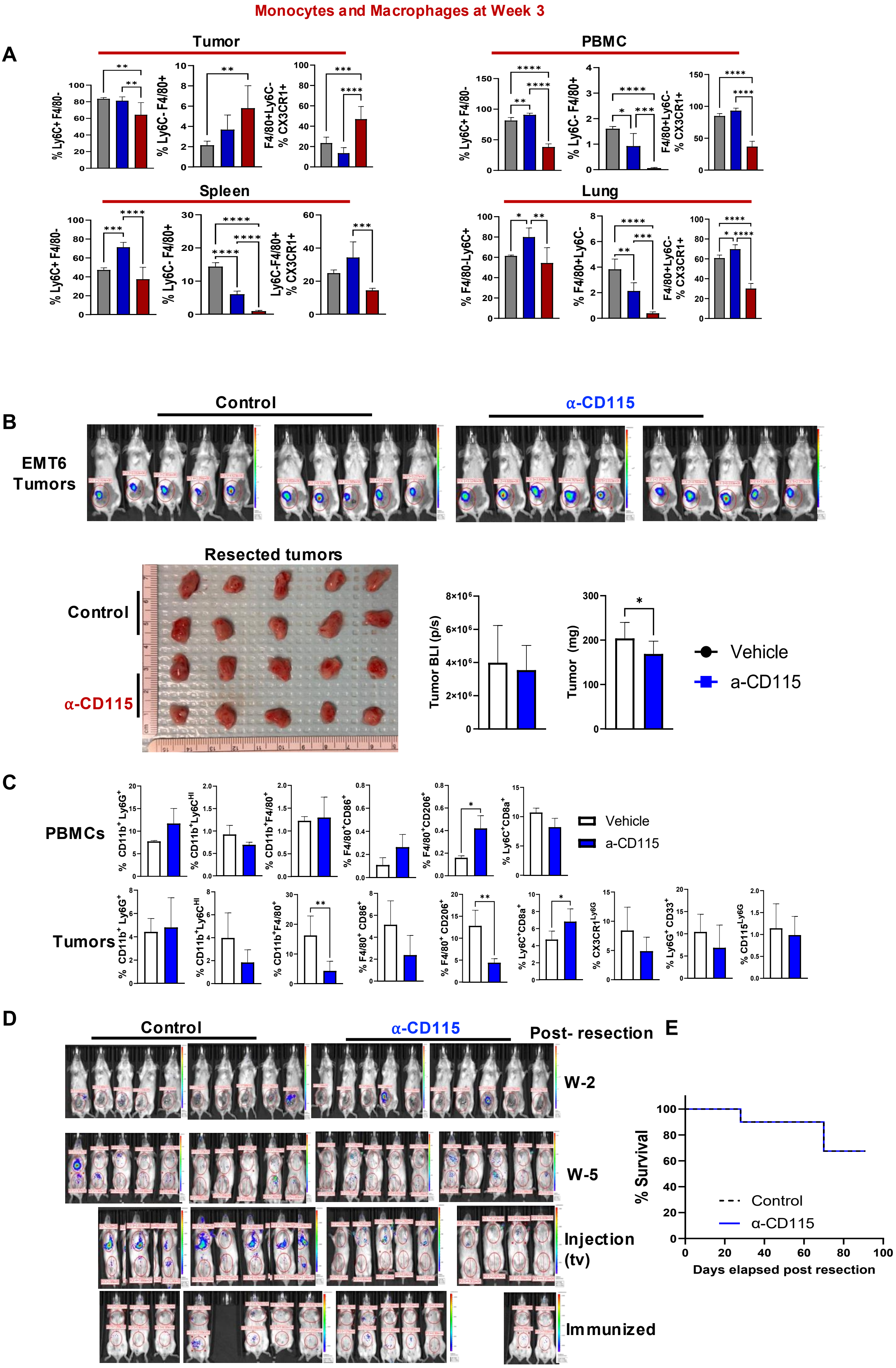


Supplemental Figure S4

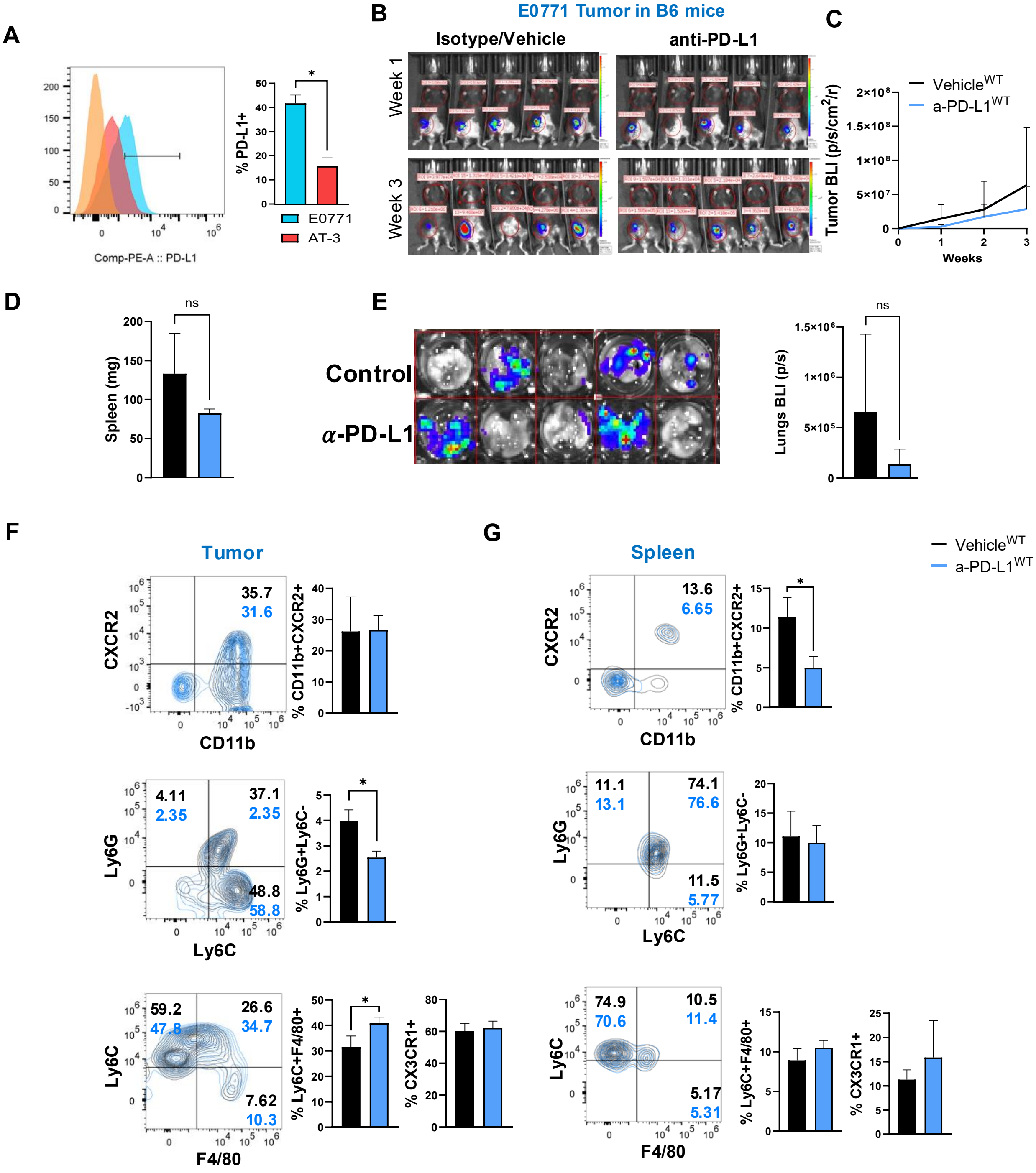


**Supplemental Figure S5**
